# Dysregulated Ca^2+^ signaling, fluid secretion, and mitochondrial function in a mouse model of early Sjögren’s syndrome

**DOI:** 10.1101/2024.03.19.585719

**Authors:** Kai-Ting Huang, Larry E. Wagner, Takahiro Takano, Xiao-Xuan Lin, Harini Bagavant, Umesh Deshmukh, David I. Yule

## Abstract

Saliva is essential for oral health. The molecular mechanisms leading to physiological fluid secretion are largely established, but factors that underlie secretory hypofunction, specifically related to the autoimmune disease Sjögren’s syndrome (SS) are not fully understood. A major conundrum is the lack of association between the severity of inflammatory immune cell infiltration within the salivary glands and glandular hypofunction. In this study, we investigated in a mouse model system, mechanisms of glandular hypofunction caused by the activation of the stimulator of interferon genes (STING) pathway. Glandular hypofunction and SS-like disease were induced by treatment with 5,6-Dimethyl-9-oxo-9H-xanthene-4-acetic acid (DMXAA), a small molecule agonist of murine STING. Contrary to our expectations, despite a significant reduction in fluid secretion in DMXAA-treated mice, *in vivo* imaging demonstrated that neural stimulation resulted in greatly enhanced spatially averaged cytosolic Ca^2+^ levels. Notably, however, the spatiotemporal characteristics of the Ca^2+^ signals were altered to signals that propagated throughout the entire cytoplasm as opposed to largely apically confined Ca^2+^ rises observed without treatment. Despite the augmented Ca^2+^ signals, muscarinic stimulation resulted in reduced activation of TMEM16a, although there were no changes in channel abundance or absolute sensitivity to Ca^2+^. However, super-resolution microscopy revealed a disruption in the intimate colocalization of Inositol 1,4,5-trisphosphate receptor Ca^2+^ release channels in relation to TMEM16a. TMEM16a channel activation was also reduced when intracellular Ca^2+^ buffering was increased. These data are consistent with altered local coupling between the channels contributing to the reduced activation of TMEM16a. Appropriate Ca^2+^ signaling is also pivotal for mitochondrial morphology and bioenergetics and secretion is an energetically expensive process. Disrupted mitochondrial morphology, a depolarized mitochondrial membrane potential, and reduced oxygen consumption rate were observed in DMXAA-treated animals compared to control animals. We report that early in SS disease, dysregulated Ca^2+^ signals lead to decreased fluid secretion and disrupted mitochondrial function contributing to salivary gland hypofunction and likely the progression of SS disease.

## Introduction

Saliva plays crucial roles in oral health, including lubricating the mouth, maintaining pH balance, defense against microorganisms, aiding taste, and initiating digestion of macronutrients (1–3). Saliva is produced primarily by three major salivary glands; the submandibular gland (SMG), parotid gland (PG), sublingual gland (SLG), and some minor glands in the lower lip, tongue, and cheek. Saliva is generated in secretory acinar cells, with its content adjusted by ducts before reaching the mouth. The acinar cells are fundamental to the production of the primary salivary secretion (4). The fluid secretion process is driven by the trans-epithelial movement of Cl^-^ across acinar cells. To accomplish vectorial movement of Cl^-^, acinar cells are polarized such that the basolateral plasma membrane (PM) faces the interstitium and is adjacent to blood vessels, while the apical PM forms a lumen with the distinct PM regions physically segregated by tight-junctional complexes. At the basolateral PM, Cl^-^ are transported into the acinar cell cytoplasm against their electrochemical gradient *via* the Na^+^/K^+^/2Cl^-^ cotransporter, (NKCC1). Following mastication or the experience of the taste and the smell of food, the neurotransmitter, acetylcholine (ACh) is released from parasympathetic nerves and acts on muscarinic receptors on the basolateral PM. Activated muscarinic receptors promote the production of inositol 1,4,5 trisphosphate (IP_3_), and subsequently Ca^2+^ release from endoplasmic reticulum (ER) stores *via* IP_3_ receptors (IP_3_Rs) situated in the ER in the extreme apical aspects of the cell (5–7). Elevated [Ca^2+^]_i_ activates a Ca^2+^-activated Cl^-^ channel named TMEM16a that allows Cl^-^ to move through the apical PM to the ductal lumen which is continuous with the salivary intercalated duct (3,8). In turn, Na^+^ moves through the paracellular space to balance the Cl^-^ and water follows osmotically both paracellularly and through the water channel aquaporin5 (AQP5) to generate the primary saliva(8).

The importance of saliva is underappreciated in the absence of hypofunction. Reduced salivary secretion is termed xerostomia and can result from the iatrogenic effects of drugs, as collateral damage to salivary glands following radiotherapy for malignancy in the head and neck area, and commonly in Sjögren’s syndrome (SS) (9). SS is a chronic autoimmune disorder, that is predominantly manifested as profound dry eye and dry mouth as ultimately the immune system targets and destroys lacrimal and salivary gland cells (10–13). SS can occur independently (primary SS, pSS) or concurrently with diseases such as arthritis or lupus (secondary SS, sSS) (14). SS affects millions of people, predominantly females in their fourth and fifth decades of life. While treatments can alleviate symptoms, there is no cure or intervention to halt its progression. The etiology of SS remains largely unresolved, but it’s believed to result from a combination of genetic, environmental, hormonal, and possibly viral factors, causing an aberrant immune response directed against the exocrine glands. The identification of SS usually is scored by the extent of salivary hypofunction, the degree of immune infiltration, evidence of damage to minor salivary glands observed following biopsy, and the presence of autoantibodies, such as anti-SSA (Ro) and Anti-SSB (La) and anti-nuclear antibody (ANA) which are classically found in SS (12,13,15). Notably, however, in the early phases of SS, there is minor immune infiltration and little overt damage to exocrine tissue despite profound hypofunction. Provocatively, these data indicate that loss of secretory tissue *per se* is not the causative event resulting in dryness early in the disease, and further indicates that a defect in the stimulus-secretion coupling mechanism precedes glandular destruction and possibly contributes to the progression of the disease.

Over the years, numerous mouse models both genetic and “induced” have been developed to study the pathogenesis of SS, with each exhibiting specific aspects of the human condition, including glandular dysfunction, autoantibody production, and lymphocytic infiltration (16,17). To investigate the early events in SS, we concentrated on an SS model induced by the activation of the stimulator of the interferon gene (STING) pathway. This is thought to mirror the molecular response to bacterial/viral infection. STING is primarily located in the endoplasmic reticulum (ER) and plays a crucial role in the innate immune response, especially against DNA viruses and intracellular bacteria. Activation of STING occurs upon sensing cytosolic DNA as a result of cell damage or from microbial origin following infection. When cytosolic DNA is detected, it is first recognized by a sensor molecule called cGAS (cyclic GMP-AMP synthase). Binding to DNA prompts cGAS to generate cGAMP (cyclic GMP-AMP), which, in turn, binds to and activates STING (18). Once STING is activated, it undergoes a series of transformations that ultimately result in the transcription of type I interferon genes, especially interferon-β (IFN-β) (19). The production of type I interferons is a primary antiviral response and a significant characteristic of SS (19,20). STING can be activated pharmacologically by exposure to 5,6-Dimethyl-9-oxo-9*H*-xanthene-4-acetic acid (DMXAA), which faithfully reproduces the immune response observed following physiological STING activation (21–23).

In this study, we investigated the early events in the initiation of SS-like disease that lead to salivary gland hypo-function using the DMXAA SS model. We first utilized *in vivo* intravital imaging to investigate any potential dysregulation of Ca^2+^ signaling in the DMXAA-induced SS mouse model. Paradoxically, the spatially averaged Ca^2+^ levels achieved following neural stimulation in mice treated with DMXAA were enhanced despite significantly reduced fluid secretion. Notably, however, the stereotypical spatial characteristics of the Ca^2+^ signal were disrupted. Downstream of the Ca^2+^ signal, the activity of the TMEM16a Ca^2+^-activated Cl channel stimulated by muscarinic secretagogues was reduced, despite no changes in the abundance or localization of the protein or absolute sensitivity to activation by Ca^2+^. The intimate localization of IP_3_R and TMEM16a was however disrupted, contributing to the reduced activity of TMEM16a upon agonist stimulation as local peripheral coupling between channels is disrupted. Moreover, we observed disordered mitochondrial morphology, abundance, and function in the disease model. These data suggest that early in SS, reduced fluid secretion occurs because of a defect in the secretagogue activation of Cl^-^ secretion. Further significant mitochondrial dysfunction is evident, possibly as a result of the aberrant Ca^2+^ signals that may contribute to the dysregulated Ca^2+^ signals and/or progression of SS disease.

## Results

### Saliva secretion is attenuated in both SMG and PG in the SS mouse model

Activation of the STING pathway in mice has been established as a model for the initiation of SS. This pathway is normally activated following exposure to foreign nucleic acids and is thought to mimic exposure of cells to DNA/RNA from viruses and bacteria. Activation of this pathway in salivary glands is characterized by initiation of a type-1 interferon response, mild immune cell infiltration, and a marked loss of saliva secretion without obvious morphological damage and therefore mimics the early clinical manifestations of SS disease. Thus, to investigate the early cellular events in acinar cells during the initiation of SS in mice, we chose to pharmacologically activate this pathway using DMXAA, a STING pathway agonist. As described in Methods, DMXAA (or control solution) was administered on day 0 and day 21 of the experimental timeline (Figure 1A). Immunofluorescent staining in sliced SMG tissue indicated that STING protein was increased in SMG in the DMXAA-treated mouse on day 28, seven days after the final DMXAA administration (supplementary Figure 1) confirming the activation of the STING pathway. Whole saliva production from the major salivary glands was evaluated on day 28 following systemic stimulation with the muscarinic receptor agonist, pilocarpine. To avoid potential weight-related variations in saliva secretion, the total saliva output was normalized to the individual mouse’s body weight. Notably, the average saliva production was reduced from 130.1± 48.96 mg in vehicle-treated mice to 63.71± 30.41 mg in DMXAA-treated mice, a reduction in saliva production of 48.97% (Figure 1B). Consequently, DMXAA treatment resulted in 51.99% saliva production compared to vehicle-treated mice (Figure 1C). Moreover, H&E staining indicated that mild immune infiltration was observed in the DMXAA-treated mice with no overt changes to the morphology of the gland (Figure 1D). The mass of the SMG was also not significantly different in DMXAA vs. vehicle-treated animals, consistent with no loss of secretory tissue (Figure 1E). Collectively, these results suggest that DMXAA-treated mice exhibit characteristics of early-stage SS and could be a useful model for investigating the pathophysiological mechanisms underlying secretory dysfunction and advancing our understanding of the disease’s progression.

**Figure 1.**
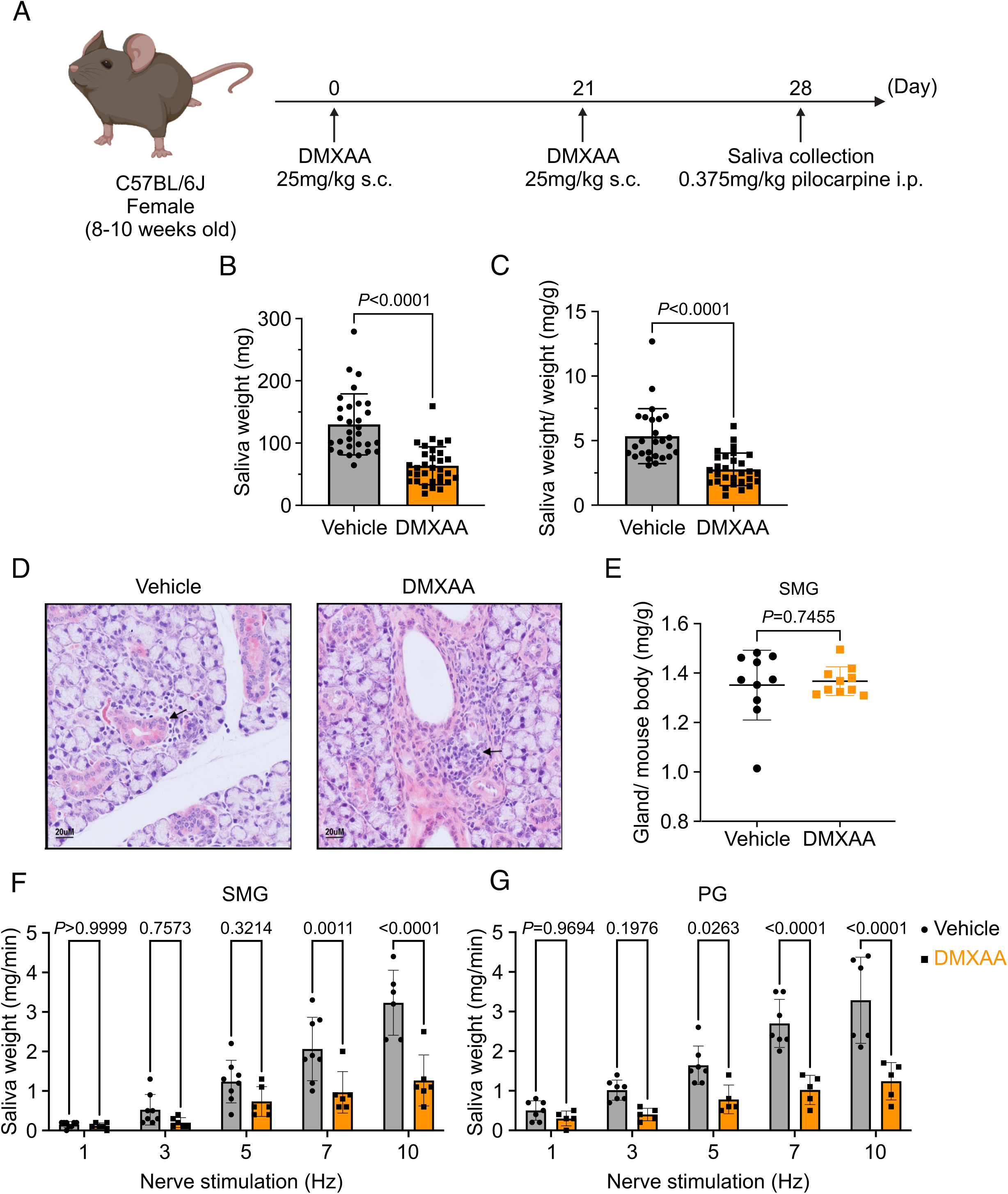
Deficiency in salivary secretion in DMXAA-induced SS mouse model. (A) Schematic timeline for the generation of the SS mouse model. Female wild-type (WT) mice were administered two subcutaneous doses of DMXAA on Day 0 and Day 21. Salivary gland function was assessed on day 28. (B-C) Saliva, stimulated by pilocarpine, was collected over 15 minutes. (B) The amount of saliva secretion was determined by measuring the saliva weight. Vehicle: N= 30 mice, SS mouse model: N= 32 mice. Mean ± SD. (C) The weight of collected saliva was normalized to each mouse’s body weight. Vehicle: N= 26 mice, SS mouse model: N= 29 mice. Mean ± SD. Unpaired two-tailed t-test. (D) H&E stained sections from the vehicle or DMXAA-treated animals. Treated animals showed minor lymphocyte infiltration and inflammation as focal peri-vascular/peri-ductal lymphocytic sialoadenitis adjacent to normal-looking acini. (E) The glandular damage was assessed by normalizing the weight of the SMG to the mouse’s body weight. Each dot represents the weight of one SMG. N=10 from 5 mice for both vehicle-treated and DMXAA-treated mice. (F-G) A comparison of total saliva secretion following 1 min stimulations at the indicated frequency (F) from the SMG of mice (Vehicle: N= 8 mice, SS mouse model: N= 6) and (G) from the PG (Vehicle: N= 7 mice, SS mouse model: N= 5 mice). Mean ± SD. Two-way ANOVA with multiple comparisons.

To further investigate the individual relative contribution of the SMG and PG to the decrease in total saliva secretion using a more physiological stimulation paradigm, we performed experiments where the nerve bundle innervating a particular gland was electrically stimulated and saliva secretion quantitated. Previous research in our lab has established the range and parameters for physiological stimulation of secretion (24). The production of saliva was significantly diminished in DMXAA-treated animals compared with vehicle controls at stimulation frequencies of 7 and 10 Hz in SMG (Figure 1F) and at 5, 7, and 10 Hz in the PG (Figure 1G). These findings confirm that activation of the STING pathway reduces the function of both SMG and PG, consistent with the diminished production of whole saliva. Notably, the reduction in function of PG, the gland responsible for the majority of stimulated saliva secretion, was relatively greater than in SMG.

### Altered spatiotemporal characteristics of Ca^2+^ signals in the SS mouse model

An increase in intracellular Ca^2+^ plays a central role in regulating the cellular machinery underlying the fluid secretion mechanism. In particular, as noted, an increase in Ca^2+^ is important for the activation of ion channels localized in particular domains of the polarized acinar cell which play a central role in saliva secretion (24). Given that the precise spatiotemporal characteristics of the Ca^2+^ signal in salivary acinar cells are thought to be fundamental to the appropriate activation of the fluid secretion machinery, we evaluated whether hypofunction following DMXAA treatment resulted from dysregulation of the stimulated Ca^2+^ signal. Previous research in our lab developed a platform to study Ca^2+^ signaling *in vivo* using Multiphoton (MP) imaging in transgenic mice engineered to express a genetic-encoded Ca^2+^ indicator, GCaMP6f, specifically in acinar cells (24). The protocol for STING pathway induction was applied to the Mist1^CreERT+/-^GCaMP6f^+/-^ genetic mouse (supplementary Figure 2A). Salivary gland function was assessed by the amount of pilocarpine-induced saliva secretion. The secretion deficiency observed in wild-type mice was recapitulated in these mice from a different genetic background (supplementary Figure 2B). We reasoned that decreased fluid secretion could result from reduced or dysregulated Ca^2+^ signaling in DMXAA-treated mice. Therefore, next, we compared the spatially averaged Ca^2+^ signal evoked by direct nerve stimulation in DMXAA-treated vs. vehicle control animals *in vivo* in SMG. SMG were stimulated at frequencies optimum for fluid secretion (1-10 Hz) for 10 seconds and the Ca^2+^ signals were recorded. Figure 2A are standard deviation (SD)-projection images visualizing the spatial and amplitude changes in Ca^2+^ throughout the field of view during the entire period of stimulation at the indicated frequencies. In vehicle-treated mice, the Ca^2+^ signals at low stimulus strengths occurred in a minority of the cells and predominantly propagated below the apical PM. As the stimulation strength increased, Ca^2+^ signals became more pronounced, and more acinar cells responded. Strikingly, acinar cells in DMXAA-treated mice demonstrated enhanced sensitivity to stimulation. At lower stimulation strengths, a larger number of acinar cells responded, and the spatially averaged Ca^2+^ signals in these cells were notably larger when compared to those in the control group. Figure 2B shows a time series of images following 7 Hz stimulation. This augmented response was manifested as an elevated maximum peak [Ca^2+^] (Figure 2D), shorter latency (Figure 2E), and larger area under the curve (AUC) during stimulation in DMXAA-treated animals (Figure 2F).

**Figure 2.**
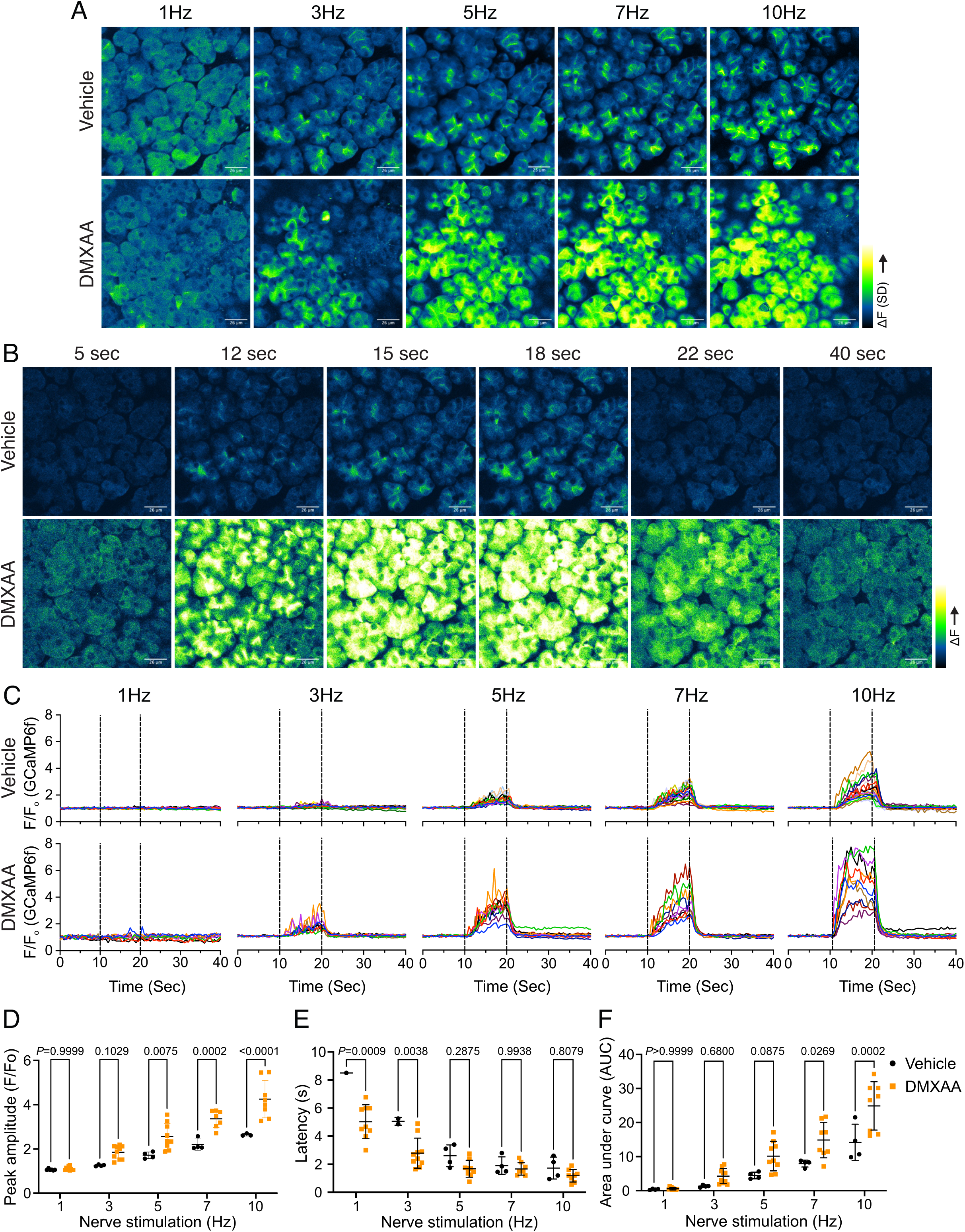
Augmented global Ca^2+^ signals *in vivo* in SS mouse model. (A) Representative standard deviation images of Ca^2+^ signals during the 10 s of stimulation. Scale bar: 26 μm (B) The time-course of pseudo-color images of Ca^2+^ in response to 7Hz stimulation. Scale bar: 26 μm (C) Representative cellular responses to stimulation at the indicated frequencies averaged from the entire cell. N = 10 cells, one animal. (D) A comparison of peak Ca^2+^, (E) area under curve, and (F) latency during each stimulation in SMG. Each symbol represented the average response of ten cells from one view. Vehicle: N= 3-6 from three mice; SS mouse model: N= 8-10 from four mice. Mean ± SD. Two-way ANOVA with multiple comparisons.

In addition to the absolute magnitude of the spatially averaged Ca^2+^ signal, the subcellular spatial characteristics of the stimulated Ca^2+^ rise are also important for appropriate stimulation of fluid secretion (24). Ca^2+^ signals stimulated following nervous stimulation in SMG are invariably initiated in the extreme apical pole of acinar cells and subsequently establish a standing gradient that dissipates rapidly to result in apically confined signals that do not substantially propagate to the basal aspects of the cell following physiological stimulation (24). We therefore investigated if the spatial characteristics of stimulated Ca^2+^ signals were altered in DMXAA-treated animals. SD image projections generated during the period of stimulation demonstrated that the [Ca^2+^]_i_ increase was tightly localized below the apical PM within the acinar cells in the vehicle-treated animals (Figure 3A). However, in the DMXAA-treated animals, the [Ca^2+^]_i_ exhibited a more global distribution through the entire cell cytoplasm (Figure 3B). The [Ca^2+^]_i_ was visualized *via* line-scan plots revealing the temporal alterations along a line extending from the apical PM to the basolateral PM, traversing the nucleus, over time within an acinar cell. A significant [Ca^2+^]_i_ elevation was evident at the basolateral aspects of the acinar cell in the DMXAA-treated animals (Figure 3D) when compared to the vehicle-treated control (Figure 3C). As shown in the kinetic plots, the [Ca^2+^]_i_ in the apical region is greater in DMXAA-treated mice compared to the vehicle-treated mice (Figure 3E). Moreover, a significant Ca^2+^ signal was observed in the extreme basal region of the cell in DMXAA but not in vehicle-treated animals (Figure 3F). The comparison of Ca^2+^ signal ratios at the apical versus basolateral PM indicated the most significant globalization of the Ca^2+^ signal occurred at 10 Hz stimulation (Figure 3G) which corresponds to the stimulation strength that results in maximal fluid secretion (24). In total, these data demonstrate that the magnitude of the spatially averaged Ca^2+^ signal, together with the spatiotemporal characteristics of the signal are altered in the DMXAA-treated animals, but that these changes can not readily account for the reduction in fluid secretion observed.

**Figure 3.**
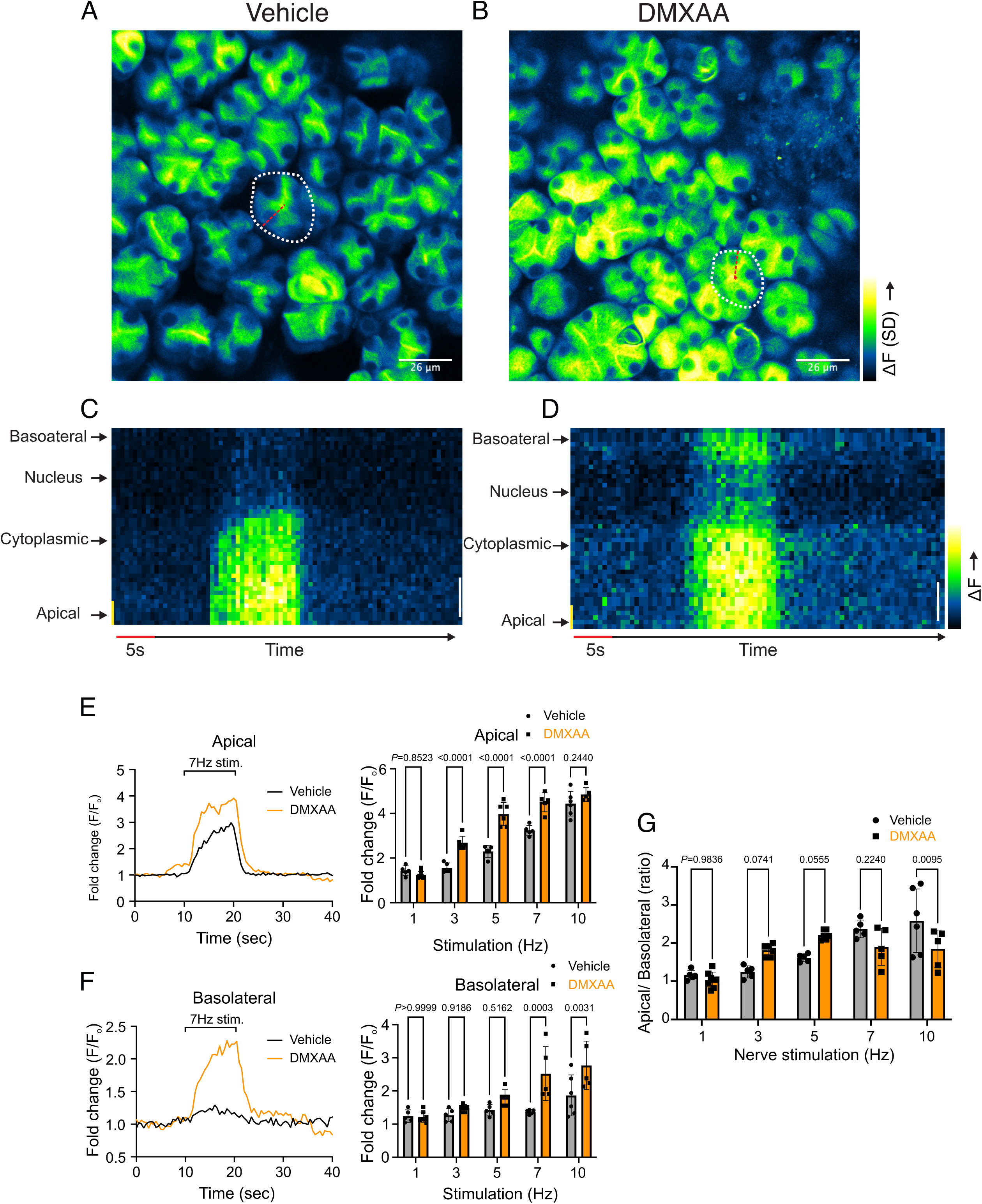
Disrupted spatial localization of Ca^2+^ signals *in vivo* in SS mouse model. (A-B) A representative standard deviation image during the 7Hz stimulation in (A) vehicle condition and (B) in the SS mouse model. Scale bar: 26 μm. An acini is outlined by the white broken line and a line from apical to basal is shown in red in each SD image. (C-D) A representative “kymograph” image of consecutive lines stacked in space over time for 7Hz stimulation in (C) vehicle condition and (D) SS mouse model. Time is encoded along the X-axis from left to right. Space is encoded along the Y-axis from the apical side (bottom) to the basolateral side (top). Scale bar: 3 μm. (E) Representative trace of Ca^2+^ signals at 7Hz nerve stimulation in an apical ROI generated as the initial 2 μm of the scanned line over time (yellow line) in vehicle-treated (black) and DMXAA-treated (orange) mice. The changes in apical ROI fluorescence at the indicated frequencies were quantified as the maximal Ca^2+^ changes normalized to the basal intensity. (F) Representative trace of Ca^2+^ signals following 7Hz nerve stimulation in a basolateral ROI generated as the final 2 μm of the scanned line (yellow line) over time in vehicle-treated (black) and DMXAA-treated (orange) mice. The changes in basolateral Ca^2+^ signals at the indicated frequencies were quantified by the maximal Ca^2+^ changes normalized to the basal intensity. (G) The ratio of the magnitude of Ca^2+^ signal on the apical vs. the basolateral ROI upon stimulation at the indicated frequencies. Vehicle: N= 5-6 replicates from three mice; SS mouse model: N=5-8 replicates from four mice. Mean ± SD. Two-way ANOVA with multiple comparisons.

### Secretagogue stimulated TMEM16a activity is suppressed in the SS mouse model

The rate-limiting step for the secretion of fluid is the activation of the Ca^2+^-activated Cl^-^ channel, TMEM16a. We considered that a reduction in fluid secretion could conceptually occur by a reduction or mislocalization of TMEM16a protein, or by compromised muscarinic receptor-stimulated activation of the channel. Western blotting indicated the TMEM16a protein expression was comparable between the vehicle and SS mouse models (Figure 4A and 4B). In addition, immunolocalization using confocal microscopy demonstrated that TMEM16a localization remained largely unchanged in the SS mouse model (Figure 4C). Thus, a decrease in protein expression or mislocalization of the protein does not result in a reduction in stimulated saliva secretion. We next investigated whether the activation of this ion channel was compromised in DMXAA-treated animals using whole-cell patch clamp electrophysiology. In the absence of stimulation, no Cl^-^ currents were observed in either vehicle or DMXAA animals following either depolarizing or hyperpolarizing voltage steps from a holding potential of −50 mV (Figure 4D). In the presence of 1 μM of muscarinic agonist Carbachol (CCh), robust Cl^-^ currents were measured in acini prepared from vehicle-treated animals (Figure 4E), which were greatly reduced in DMXAA-treated animals (Figure 4D and 4F). Reduced CCh-stimulated Cl^-^ currents could potentially occur because of altered Ca^2+^ regulation of TMEM16a following disruption of the spatial characteristics of the stimulated Ca^2+^ signal. Theoretically, it is also possible that the [Ca^2+^]_i_ in the immediate vicinity of TMEM16a was disrupted, despite the augmented spatially averaged peak response. We therefore next tested whether TMEM16a activity stimulated directly by 0.5, 1 or 5 μM Ca^2+^ in the pipette solution (and thus globally in the cytoplasm) was altered in DMXAA-treated animals. Surprisingly, TMEM16a was activated to a similar extent by Ca^2+^ in the SS mouse model (Figure 5A and 5B). In total, our data suggest that TMEM16a abundance, localization, or activity *per se* are not altered and thus do not explain the significant reduction in saliva secretion in the DMXAA-treated model.

**Figure 4.**
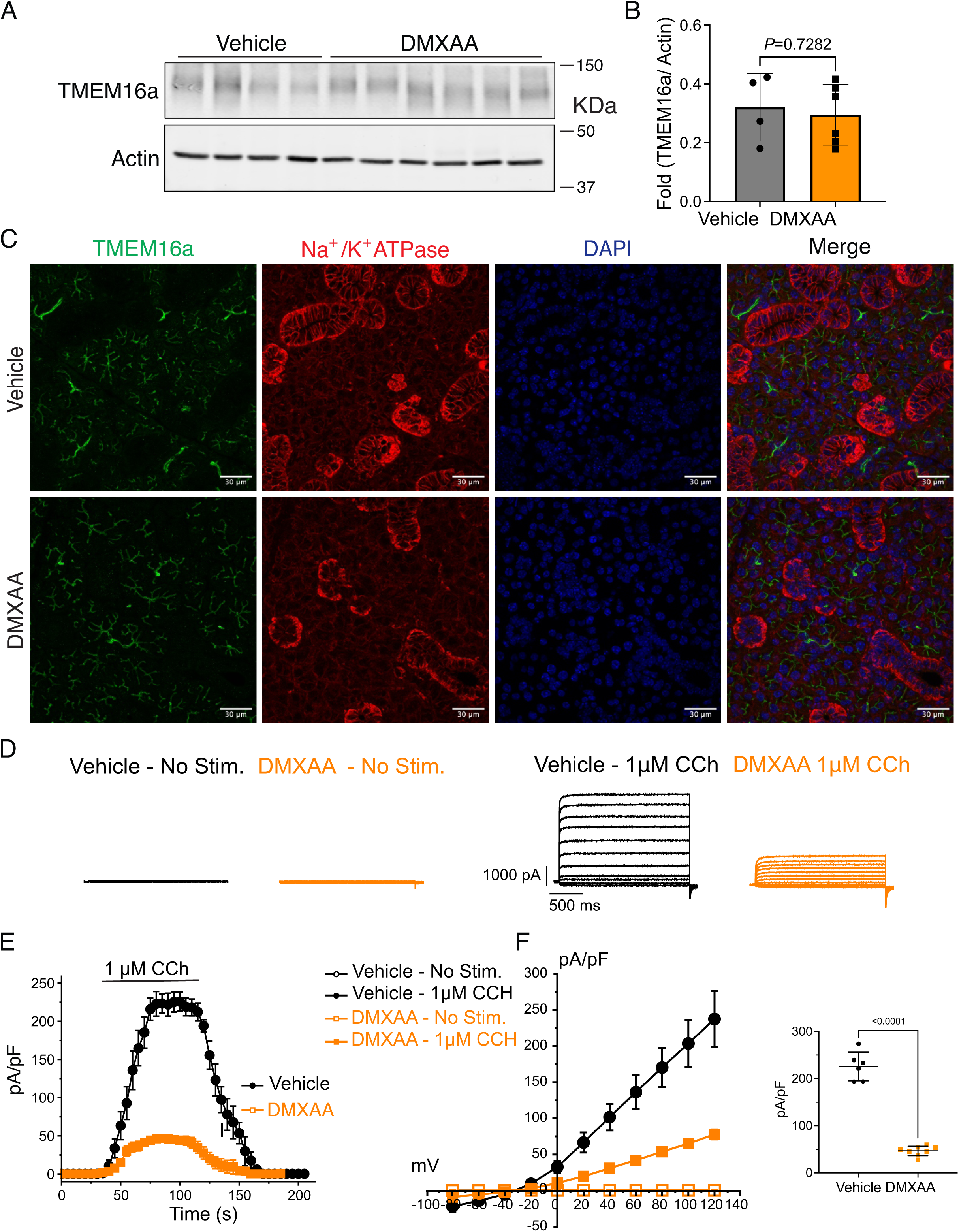
Attenuated whole-cell macroscopic Cl^−^ currents induced by CCh stimulation in SS mouse model. (A) Western blotting showing the protein expression level of TMEM16a in the vehicle condition and the DMXAA-treated SS mouse model. Actin is the internal control. (B) The quantification of TMEM16a protein expression normalized to the internal control, Actin. Vehicle, N= 4 mice; SS mouse model: N= 6 mice. (C) Immunofluorescent staining in SMG tissue for TMEM16a (green), Na^+^/K^+^ ATPase (red), and DAPI for nucleus (blue). The upper panel is from the vehicle-treated control and the bottom panel is from DMXAA-treated animals. Scale bar: 30 μm. Unpaired two-tailed t-test. (D) Cl-currents when cells were held at −50 mV and stepped from −80 to 120 mV in 20 mV increments. (E) Time-dependent Cl^-^ current density changes in response to the CCh in the isolated acinar cells in vehicle conditions and SS mouse model. (F) Current-voltage relationships were measured before and after the addition of CCh in vehicle conditions (N=three mice, 3-4 cells per mouse) and SS mouse model (N=three mice, 3-4 cells per mouse). TMEM16a currents in the treated mice were markedly reduced compared to the control mice. Black dots represent the vehicle-treated cells and orange squares represent DMXAA-treated cells. The open symbols represent no stimulation; the solid symbols represent CCh stimulation.

**Figure 5.**
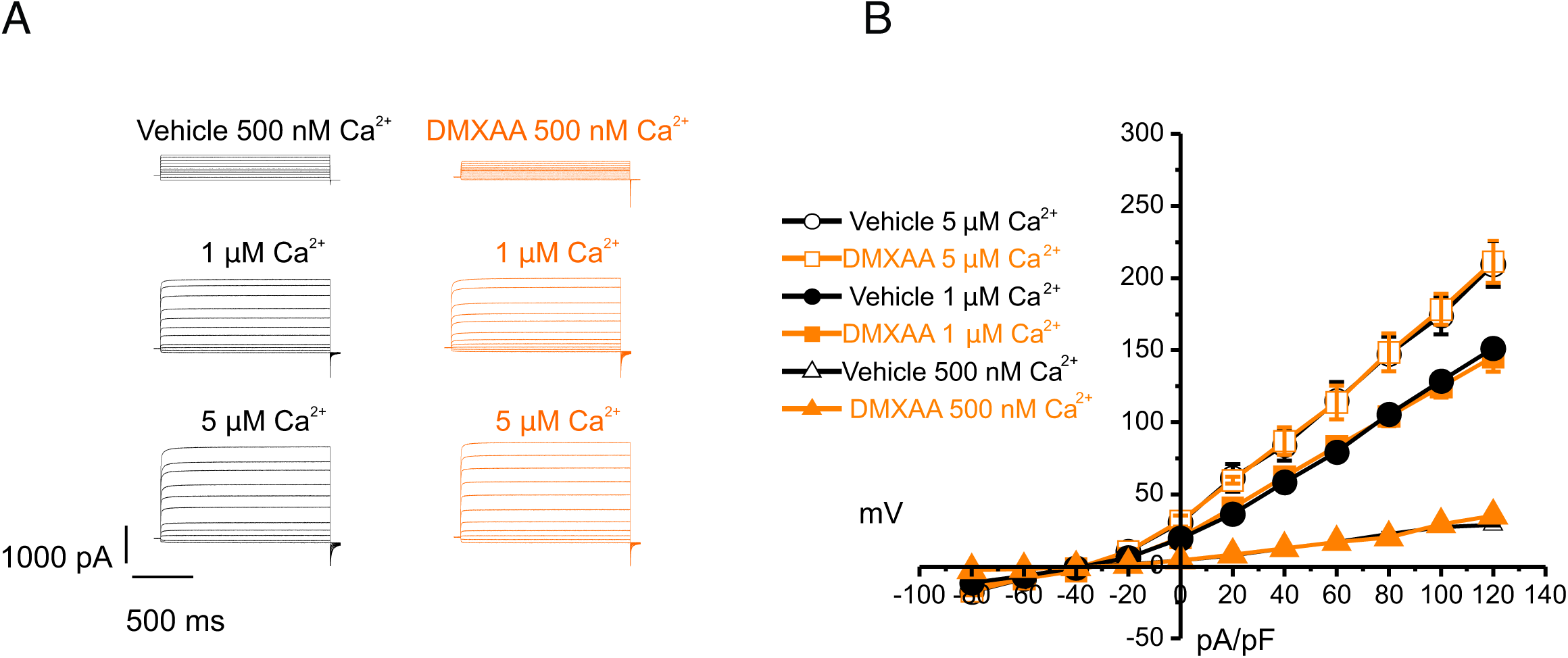
Increased [Ca^2+^]_i_ is capable of restoring TMEM16a functionality to DMXAA-treated mice. (A) Cl^-^ currents when cells were held at −50 mV and stepped from −80 to 120 mV in 20 mV increments. Either 0.5, 1 or 5 μM [Ca^2+^]_i_ in the patch pipette elicited a similar magnitude of Cl^-^ currents for both the treated (N= 3 mice, 3-4 cells per mouse) and control mice (N= 3 mice, 3-4 cells per mouse). (B) Current-voltage relationships for both populations were essentially identical. Vehicle and SS mouse model: N=3 mice, 3-4 cells per mouse.

An alternative mechanism could be that the microdomain between the apical ER Ca^2+^ release sites and the apical PM TMEM16a is disrupted in the disease model, resulting in compromised local coupling between the ER and PM channels. Therefore, we investigated how the activation TMEM16a was affected by buffering the CCh-stimulated cytosolic Ca^2+^ with slow and fast Ca^2+^ chelators. EGTA is a high-affinity Ca^2+^ buffer with slow kinetics, while BAPTA has much more rapid kinetics. Experimentally, BAPTA can quickly buffer Ca^2+^ changes close to Ca^2+^ release sites and limit local activation of effectors within 20 nm. In contrast, EGTA has been shown to attenuate the rise in the bulk cytosol but is too slow to buffer local Ca^2+^ in a restricted microdomain (25). In the presence of BAPTA, no CI^-^ currents were detected in either vehicle-treated or DMXAA-treated animals (Figure 6A). However, in the presence of the slow Ca^2+^ chelator, EGTA, CCH-stimulated Cl^-^ currents were observed in acinar cells from vehicle-treated animals but were absent in the DMXAA-treated animals (Figure 6A and 6B). These data suggest that physiologically TMEM16a is activated by local changes in Ca^2+^ signaling as a result of IP_3_-induced Ca^2+^ release in the apical domain of the acinar cell.

**Figure 6.**
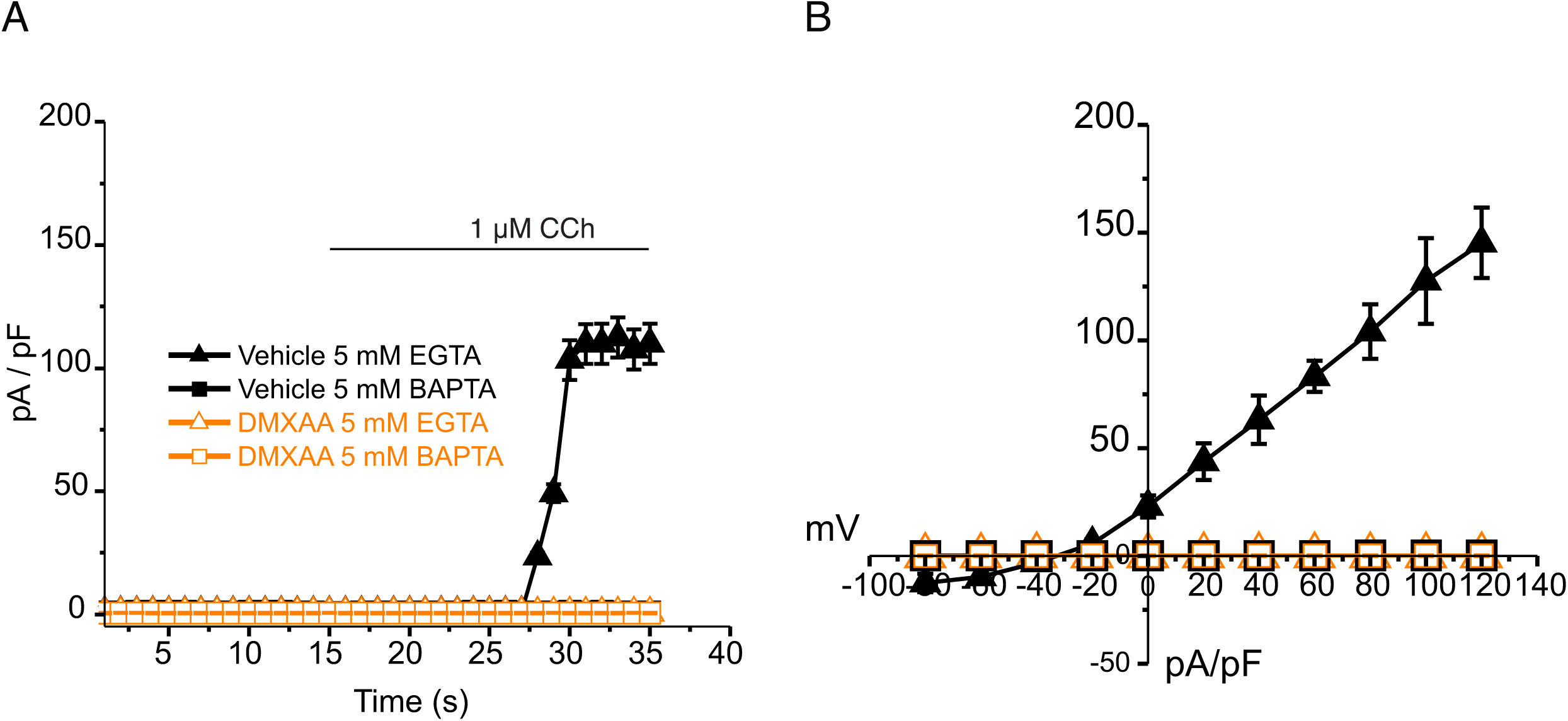
EGTA abolishes TMEM16a currents in DMXAA-treated mice. (A) Cl^-^ currents in cells held at −80 mV and stepped to 80 mV with CCh addition in EGTA (slow) and BAPTA (fast) buffered cells, respectively. (B) Current-voltage relationships were measured after the addition of CCh in 5 mM EGTA and 5m M BAPTA loaded isolated acinar cells from vehicle conditions (N= 3 mice, 3-4 cells per mouse) and SS mouse model (N= 3 mice, 3-4 cells per mouse). No TMEM16a currents in acini in either vehicle or DMXAA-treated mice in cells buffered with BAPTA. Triangles represent the 5 mM EGTA condition; squares represent the 5 mM BAPTA condition. The solid black symbols represent the vehicle-treated cells and hollow orange symbols represent DMXAA-treated cells.

Next, we employed STED super-resolution microscopy to closely examine the spatial relationship between apical PM TMEM16a and IP_3_R3 on the apical ER (Figure 7A). Despite the cell-cell contact distance remaining consistent in the disease model, as indicated by the distance between TMEM16a on the PM of adjacent acinar cells (Figure 7F), a notable increase in distance between the apical TMEM16a and IP_3_R3 expressed on apical ER compared to the control group was observed (Figure 7E). In the control mice, the distance between TMEM16a and IP_3_R3 was on average 84 ± 17 nm, versus 155 ± 20 nm in the SS disease mice. Similarly, the distance between IP_3_R3 in adjacent cells was increased from 505 ± 34 nm to 689 ± 68 nm (Figure 7C and 7D). In total, these observations support the conclusion that the reduced activity of the TMEM16a channel is attributable to the disruption of the microdomain between TMEM16a and IP_3_R3, such that the Ca^2+^ flux through the IP_3_R is not communicated appropriately to its effector, TMEM16a.

**Figure 7.**
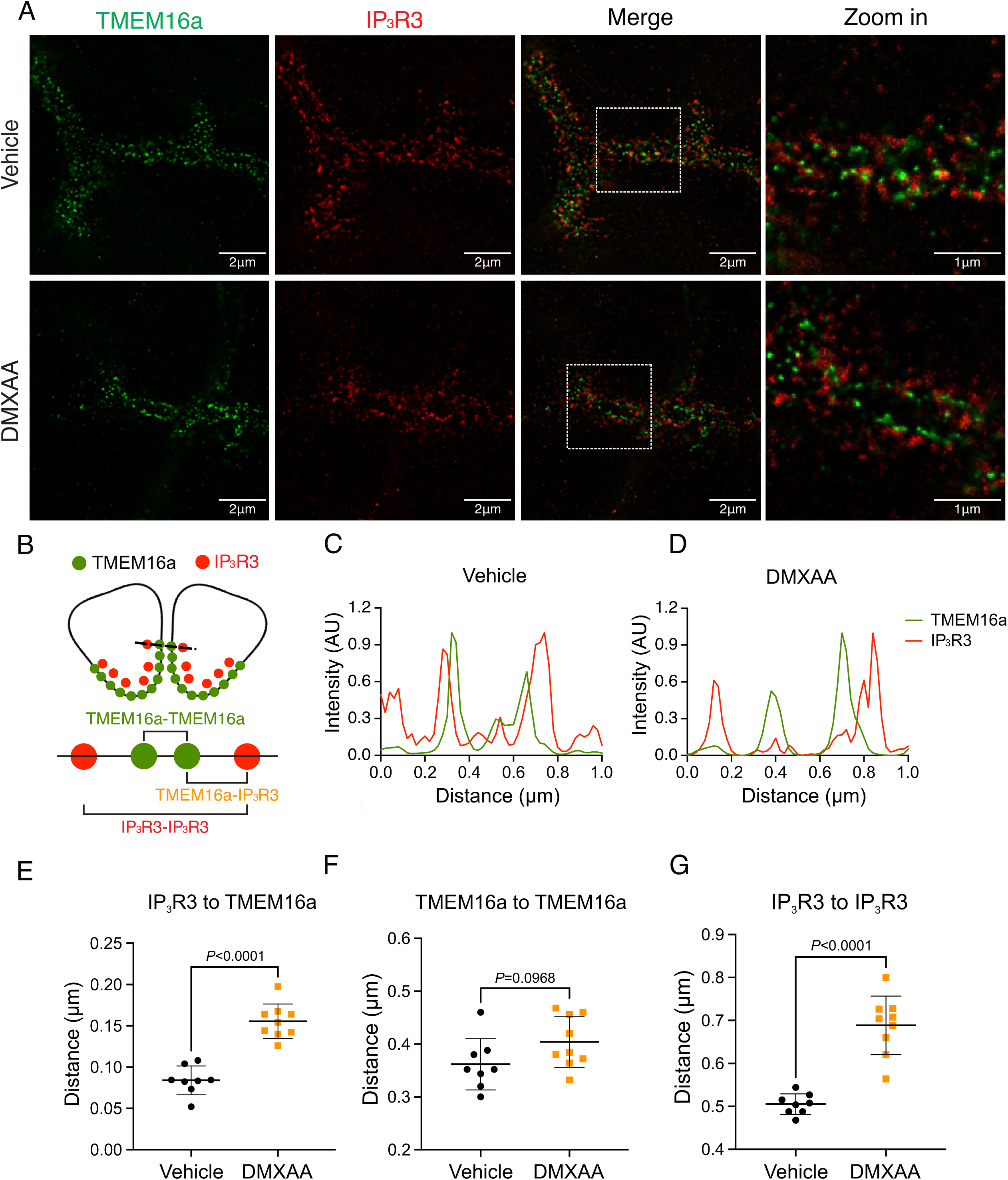
Disrupted proximity between TMEM16a and IP_3_R3 in the DMXAA-treated SS mouse model. (A) Maximum projection of a STED z stack (1 μm) showing TMEM16a (green) and IP_3_R3 (red) in SMG tissue following Huygens deconvolution. The top panel represents the vehicle-treated control, and the bottom panel represents the SS mouse model. Scale bar: 2 μm. Zoomed images highlight the localization of TMEM16a and IP_3_R3 from the white square on the merged images. (B) Diagram illustrating the positioning of apical PM TMEM16a and apical IP_3_R3 in acinar cells. To analyze the proximity, a 1 μm reference line was drawn across the two parallel TMEM16a over two adjacent acinar cells with IP_3_R3 aligned vertically in the cytoplasm. (C-D) The representative traces of changes in fluorescence of TMEM16a (green) and IP_3_R3 (red) over the 1μm distance. (E) Analysis of distance between TMEM16a and IP_3_R3 within cells. (F) Analysis of the distance between parallel TMEM16a on adjacent acinar cells. (G) Distance measurement of apical IP_3_R3 between two cells. Each symbol represents the mean of 5 examinations per image. Vehicle: N= 8 replicates from 3 mice; SS mouse model: N= 9 replicates from 3 mice. Mean ± SD. Unpaired two-tailed t-test.

### Compromised mitochondrial morphology and metabolism in the SS mouse model

Ca^2+^ modulates cellular metabolism by the intricate bidirectional interaction between the ER and mitochondria. Ca^2+^ transfer between ER and mitochondria is essential for optimal bioenergetics, and dysregulated [Ca^2+^]_i_ can be deleterious to mitochondrial function and alter morphology (26–29). The transfer of Ca^2+^ between ER and mitochondria is dependent on the intimate physical localization of the organelles (29). Notably, aberrant mitochondrial morphology has been reported in the salivary glands of SS patients (30). We first investigated mitochondrial abundance and morphology by immunofluorescence staining with antibodies directed against ATP5A, a component of the ATP synthesis machinery to visualize mitochondria, and Na^+^/K^+^ ATPase to localize the plasma membrane (Figure 8A). Using previously published methodologies (31,32), quantification revealed a 22.16% ± 4.95 reduction in mitochondrial numbers in the SS mouse model relative to the vehicle-treated control (Figure 8B). Consistent with reduced mitochondrial numbers, less area was occupied by mitochondria in DMXAA-treated acinar cells (Figure 8C). Mitochondria morphology is intricately linked to their bioenergetic status We next evaluated mitochondrial morphology by their “so-called” aspect ratio (AR) and form factor (FF) in DMXAA and vehicle-treated animals. The AR, the length of the major over minor axes of mitochondria documents the degree of fragmentation or elongation of individual mitochondria (29,32). Mitochondria exhibited an 18.35% ± 4.62 decrease in mitochondrial elongation (Figure 8D) and a 20.7% ± 7.78 decrease in mitochondrial branching (Figure 7E) in the disease model compared to the vehicle-treated control condition. Importantly, these changes in mitochondrial number and morphology were not exclusive to the SMG as similar patterns were observed in the PG mitochondria, again marked by reduced mitochondrial count, increased fragmentation, and decreased branching (supplementary Figure 3A-3E).

**Figure 8.**
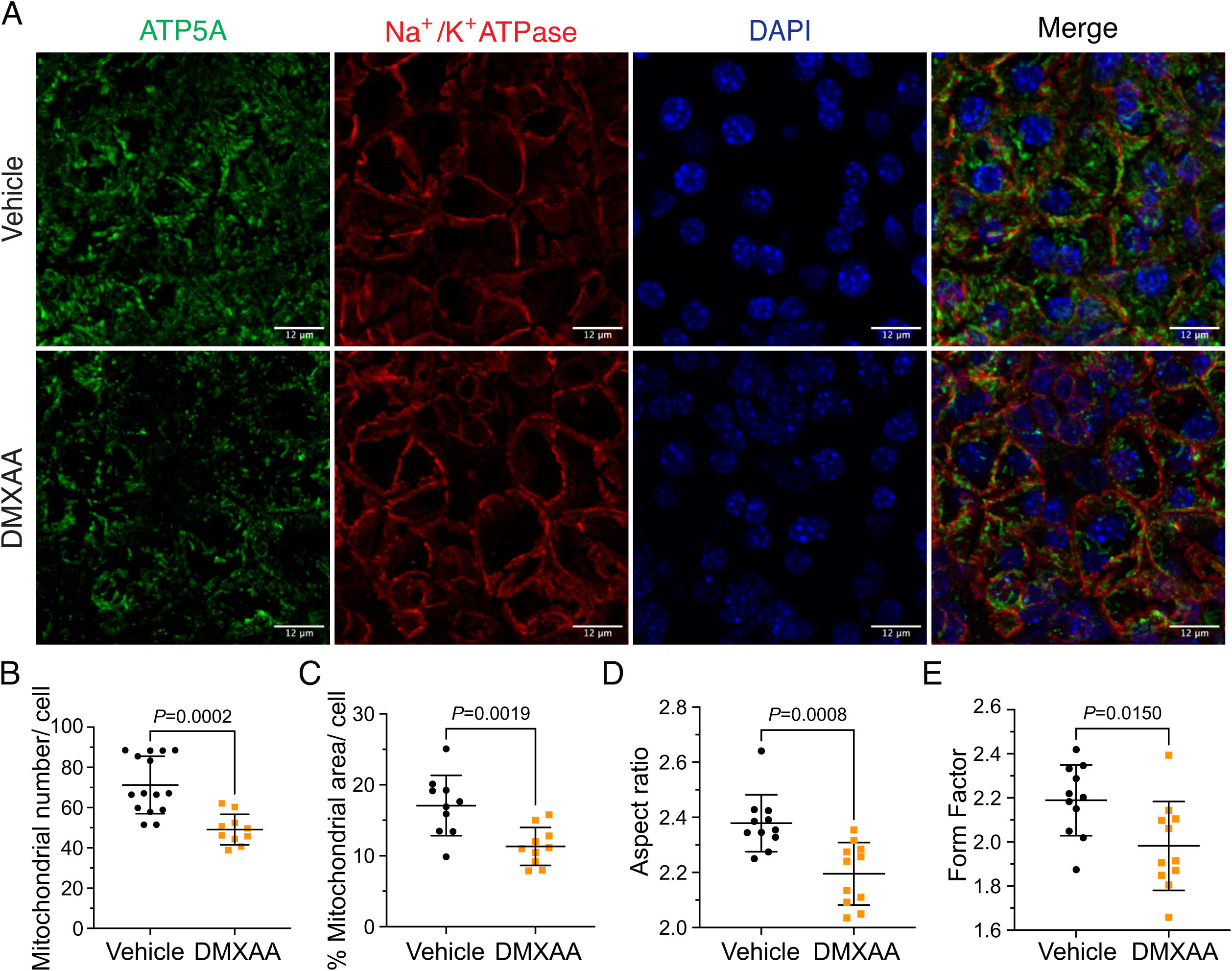
Mitochondrial alterations in acinar cells from the DMXAA-treated SS mouse model. (A) Immunofluorescent staining in SMG tissue for ATP5A (green), Na^+^/K^+^ ATPase (red), and DAPI for nucleus (blue). The upper panel is the vehicle, and the bottom panel is the SS mouse model. Scale bar: 12 μm. The mitochondrial content was quantified by (B) the mitochondrial number per acinar cell and (C) the percentage of area occupied by mitochondria per acinar cell. The mitochondrial morphology was analyzed by the (D) AR for the degree of mitochondrial tubular shape and (E) FF for the degree of mitochondrial branching (complexity). In (B) to (E), black dots represent the vehicle condition, and orange squares indicate the SS mouse model. Each symbol represents the mean of 10 cells per image. Vehicle: N= 10-15 from 3 mice; SS mouse model: N= 10-11 from 3 mice. Mean ± SD. Unpaired two-tailed t-test.

Next, we utilized electron microscopy (EM) to investigate mitochondrial ultrastructure. At low magnification, acinar cells from control mice contained defined mitochondria and well-formed ER stacks (Figure 9A. blue arrow). In contrast, the ER structure was disrupted in the SS disease model (Figure 9A’). At higher magnification, the coordinated ER structure was largely absent in diseased mice (Figure 9B and 9B’), and the proximity between ER and mitochondria was disrupted (Figure 9H and 9I). Moreover, we also observed scattered mitochondrial cristae at the highest magnification (Figure 9C’ and 9G). Consistent with immunofluorescence studies, quantification of EM micrographs revealed that mitochondria were smaller, more fragmented (Figure 9D and 9E), and rounder (Figure 9F) in shape. In summary, our results collectively indicate significant morphological alterations in mitochondria in the SS disease model.

**Figure 9.**
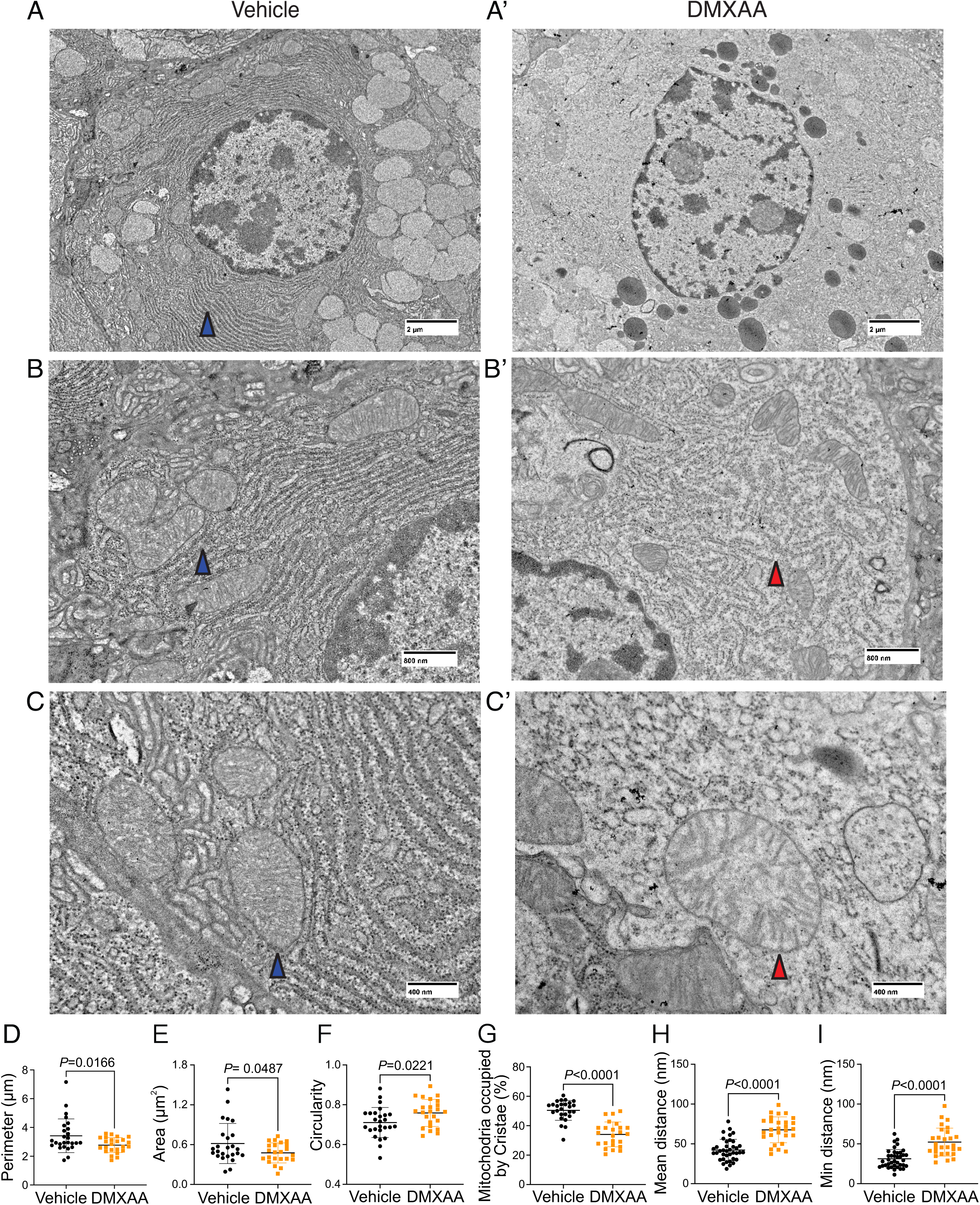
Ultrastructural analysis of mitochondria and ER in SS mouse model. (A-C’’) Images show mitochondrial cristae and ER structure by an EM at scales of (A-A’) 2μm, (B-B’) 800nm, and (C-C’) 400nm. (D) Mitochondrial perimeter, (E) mitochondrial area and (F) circularity were quantified by the shape description in ImageJ. (G) Quantification of mitochondrial cristae dispersion was evaluated by the percentage of cristae occupied in one mitochondrion. The (H) mean and (I) minimum proximity of ER and mitochondria were quantified by the plugin from http://sites.imagej.net/MitoCare/ in ImageJ. Vehicle: N=38 and SS mouse model: N=36 from 3 mice. Mean ± SD. Unpaired two-tailed t-test.

Mitochondrial morphology is a dynamic process that is intimately associated with mitochondrial bioenergetics and alterations in both, occur in response to changes in cellular status (33–36).Therefore, we investigated if changes in morphology might be associated with the disrupted function of mitochondria in the disease model. We measured mitochondrial membrane potential (ΔΨ_m_), established by the electrochemical H^+^ gradient, which is the driving force of ATP production. Isolated SMG acinar cells were loaded with TMRE, a ΔΨ_m_-specific dye, and MitoTracker Green, to confirm mitochondrial localization and to facilitate the normalization of indicator loading. The maximal z-stacks projection images taken by confocal microscopy revealed colocalization of TMRE with MitoTracker Green (Figure 10A). The basal TMRE fluorescence was reduced in cells from DMXAA vs. vehicle-treated animals (supplemental Figure 4). To assess ΔΨ_m_, we quantified the relative maximum dissipation of ΔΨ_m_ in DMXAA and vehicle-treated acinar cells by the mitochondrial uncoupler, FCCP (Figure 10B). Consistent with the reduction in basal TMRE fluorescence, the change in TMRE fluorescence normalized to mitochondrial content revealed a marked reduction in ΔΨ_m_ in the acinar cells from the SS disease model (Figure 10C).

**Figure 10.**
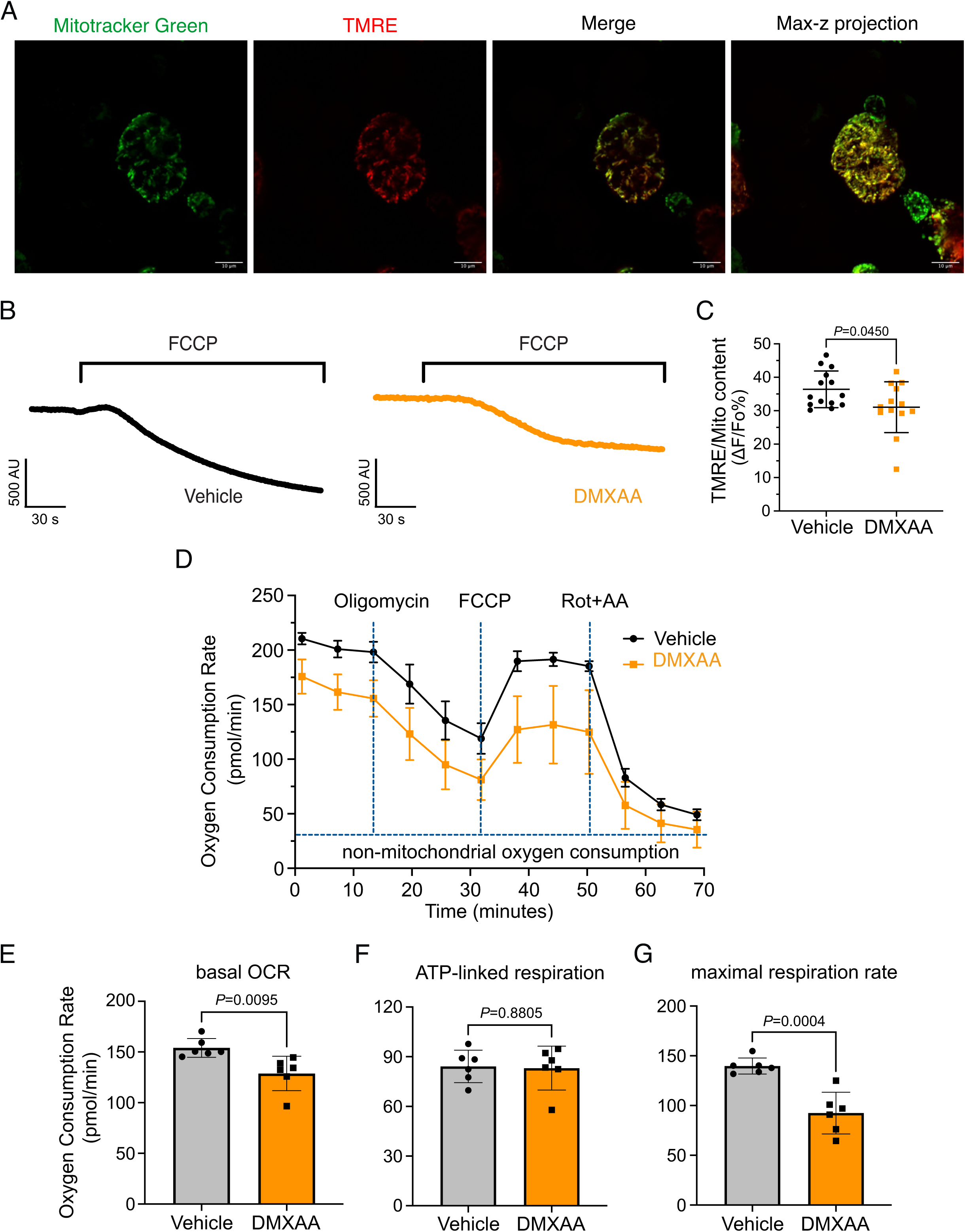
Mitochondrial bioenergetics are compromised in the DMXAA-treated SS model. (A) Mitochondria in the isolated acinar cells were labeled by the MitoTracker Green and co-stained with mitochondrial membrane potential dye, TMRE (red). The merged image shows the colocalization of both dyes, with maximal z-stack projection throughout the acinar cells. (B) Representative changes in mitochondrial membrane potential following FCCP-induced depolarization. The vehicle is shown in black; SS mouse model is in orange. (C) The quantification was achieved by the difference of TMRE normalized to MitoTracker Green. Each dot is the mean of 10 cells from one experiment. Vehicle: N=14 and SS mouse model: N=13 from 3 mice. (D) Real-time mitochondrial respiration function was assessed in isolated acinar cells from the vehicle (black) and SS mouse model (orange) using the Seahorse XFe96 extracellular flux analyzer, in response to the pharmacological mito stress (oligomycin, FCCP, rotenone, and antimycin). Vehicle: N=59 and SS mouse model: N=32 from 6 mice. (E-G) Mitochondrial respiration function parameters were quantified by OCR substracted the non-mitochondrial OCR for (E) basal respiration rate, (F) ATP-linked respiration rate, and (G) maximal respiration rate. Mean ± SD. Unpaired two-tailed t-test.

An appropriate ΔΨ_m_ mitochondrial membrane potential is vital for maintaining bioenergetics (37). Given that mitochondrial ΔΨ_m_ was significantly depolarized in DMXAA-treated animals, we next evaluated the OCR, a key metric of mitochondrial bioenergetic function in isolated SMG acinar cells. We employed sequential exposure to agents that target the function of the mitochondrial electron transport chain (ETC) using Seahorse technology (Figure 10D). Our results revealed a 25% reduction in basal OCR in the SS model compared to the control animals (at −25.25 ± 7.89 pmol/min; Figure 10E). While ATP-linked respiration showed no significant difference in post-oligomycin-induced ETC Complex V blockade in both conditions (Figure 10F). Intriguingly, the FCCP-provoked maximal respiration rate, an indicator of stress tolerance, remarkably declined by 47% ± 9.19 after FCCP treatment in the SS model (Figure 10G). These data indicate impaired mitochondrial function and stress responses in the SS mouse model.

## Discussion

SS is a complex inflammatory disease resulting from the intersection of genetics and environmental factors. This autoimmune disorder affects exocrine glands including salivary and lacrimal glands, leading to dry mouth and dry eyes, among other symptoms (11–13). SS animal models are crucial for understanding the pathogenesis, progression, and potential treatments for the disease, though like many animal models of disease, none can recapitulate all the aspects of SS. Currently, SS animal models are categorized as either those derived from genetically modified mice (14,38–40) or those where disease is induced by specific agents or environmental factors. In the context of SS, DMXAA-induced SS can be used to mimic the early stages of the disease which might be triggered in response to bacterial or viral infection. This model is particularly effective in simulating type-1 interferon immune responses seen in early SS, which is thought to contribute to the initial glandular inflammation (10,19,41). It should also be noted that DMXAA also has been reported to inhibit NAD(P)H quinone oxireductase (42) and thus the potential increase in free radical load in cells could contribute to the phenotype. The rapid symptom manifestation of disease in the DMXAA-induced model offers an advantage for investigating the early development of SS disease since DMXAA induction is a temporally controlled process, allowing the precise staging of disease onset, thus facilitating studies on the initiating events and ultimately potential early intervention and prevention strategies.

Our studies investigated stimulus-secretion coupling when fluid secretion from SMG and PG in response to physiological stimulation was significantly reduced. Previous work has established a crucial link between an increase in [Ca^2+^]_i_ and stimulation of fluid secretion in the salivary glands (8,24). Efficient secretion is reliant on the specific spatiotemporal regulation of secretagogue-stimulated [Ca^2+^]_i_ signals. Given this idea, our initial hypothesis was that a deficiency in secretion after DMXAA administration could be due to reduced or disrupted secretagogue-stimulated [Ca^2+^]_i_ signals. Indeed, previous work has revealed that in human SS patient acinar cells and the IL14α knock-in transgenic SS mouse model, CCh-induced [Ca^2+^] signals were diminished. This reduction was attributed to lower expression levels of the IP_3_R2 and IP_3_R3 proteins (43). To probe this hypothesis, we employed transgenic Mist1^CreERT2+/–^ x GCaMP6f^+/–^ that expresses Ca^2+^ indicator-GCaMP6f specifically in the acinar cells. Firstly, we validated that the activation of the STING pathway leads to similar salivary gland hypofunction in this genetic background (supplementary Figure 2). Surprisingly, however, DMXAA treatment led to a striking increase in the magnitude of neurally-induced spatially averaged [Ca^2+^]_i_ signals. This observation is not consistent with the loss of IP_3_R proteins being responsible for reduced fluid secretion previously reported in other SS models. Indeed, the expression of IP_3_R proteins was unchanged following DMXAA treatment (supplementary Figure 6). The discrepancy could be attributed to the stage of SS disease represented by the previous studies, with our data presenting an earlier initiating phase of SS disease prior to progression, at a time point before any notable decrease in IP_3_R proteins has occurred. The molecular mechanism responsible for augmented global Ca^2+^ signals following DMXAA treatment requires further study. Increased Ca^2+^ release/influx, or conversely reduced Ca^2+^ clearance might be responsible. It is tempting to speculate that reduced mitochondrial Ca^2+^ uptake and/or reduced PMCA and SERCA activity as a result of decreased ATP levels may contribute to enhanced cytosolic signals. Nevertheless, we suggest that the augmented Ca^2+^ signals might represent a compensatory mechanism to drive fluid secretion in the face of compromised physiological stimulus-secretion coupling. Although the Ca^2+^ signals were not reduced, the spatiotemporal characteristics of the Ca^2+^ signal were markedly disrupted. Specifically, during neural stimulation, while in control animals there is a pronounced standing gradient of [Ca^2+^] such that the [Ca^2+^] is much greater in the apical *vs.* basal aspects of the cell, in DMXAA-treated animals this gradient is largely absent as large changes in Ca^2+^ are propagated to the basal regions of the cells. It is conceivable that the alteration in magnitude coupled with changes in the spatial characteristics of the Ca^2+^ signal contributes to both the defect in fluid secretion and downstream cellular changes including mitochondrial damage to ultimately result in the progression of disease.

We investigated whether changes in the secretory machinery *per se* were altered in DMXAA-treated animals to result in hyposecretion. Salivary gland fluid secretion is dependent on TMEM16a facilitating CI^-^ flux across the apical PM as the driving force for water transport paracellularly and through AGP5 (44,45). The loss of either TMEM16a or AQP5 results in markedly attenuated fluid secretion (45–48). These findings indicate that alteration in expression level, localization, or regulation of these channels could potentially impact fluid secretion. Notably, in DMXAA-treated mice, the AQP5 expression and localization remain unchanged (supplementary Figure 5), consistent with a study in human labial minor salivary glands (49). We next examined whether the TMEM16a channel function was compromised in the model. Our electrophysiological analysis revealed a significant decrease in TMEM16a activity following CCh-induced stimulation. Again, this reduced activity was not the result of overt mislocalization or lower expression levels of the protein (Figure 4A and 4C). Interestingly, although the secretagogue-stimulated TMEM16a was reduced in acinar cells from DMXAA-treated animals, the sensitivity of the channel to direct activation by Ca^2+^ in the patch pipette appeared unaffected. IP_3_R3 Ca^2+^ release channels on the ER are located approximately 50-100 nm from TMEM16a on the PM (7). In this microdomain, confocal microscopy cannot easily distinguish the distinct localization of TMEM16a/IP_3_R, despite their localization on different membranes. However, STED super-resolution microscopy provides a much higher spatial resolution, achieving 20-80 nm to enable the differentiation of proteins within 20-80 nm of each other. Data using STED microscopy, suggest that the microdomain between apical ER IP_3_R3 and apical PM TMEM16a is disrupted in the disease model. The severe fragmentation of ER observed in EM images from DMXAA-treated animals also is consistent with an alteration in the relationship between ER and other intracellular domains. The disruption of the relative localization of these channels could conceivably result in diminished TMEM16a activity, if activation is dependent on the local [Ca^2+^] in its vicinity. Our data showing that the slow Ca^2+^ buffer EGTA eliminates TMEM16a activation in the disease model, but that currents can still be evoked in vehicle-treated controls is consistent with the activation of TMEM16a by the local Ca^2+^ signal surrounding the channel rather than the global cytoplasmic Ca^2+^ signal (44,50,51). Thus, the disruption of this apical microdomain likely alters the local Ca^2+^ signal that TMEM16a experiences leading to reduced activation and fluid secretion.

While changes in cytosolic [Ca^2+^] are vitally important for stimulating ion flux and hence fluid secretion, Ca^2+^ is also critical for numerous other physiological processes in salivary gland acinar cells. We focused on the potential effects of the dysregulated Ca^2+^ signal on mitochondrial morphology and function. Secretion is an energy-demanding process, necessitating a constant supply of ATP for numerous functions, including vesicle transport, protein modification, membrane fusion, and maintaining ion gradients. For example, the Na^+^/K^+^ ATPase pump generates the Na^+^ gradient, driving Cl^-^ transport into the cytosol of acinar cells through NKCC1, and SERCA pumps replenish ER Ca^2+^ levels. In this context, mitochondria are essential as they provide ATP, regulate Ca^2+^ homeostasis, supply metabolic intermediates, and coordinate with the ER to orchestrate cellular functions (52,53). Notably, recent studies have highlighted that mitochondria are abundant and display varied positioning and dynamics in salivary gland cells (54). In SS patients, there are notable alterations in mitochondrial structure, including swelling and disrupted cristae (30,55). Correspondingly, mitochondrial-related genes, particularly those involved in metabolism, dynamics, and the electron transport complex, are significantly affected (55). Our data, employing fluorescent immunostaining and EM, mirrors these findings in DMXAA-treated animals. We observed that mitochondrial morphology is altered such that mitochondria are more swollen and rounded, with dispersed cristae, similar to that reported in human SS patients (30). Since optimal mitochondrial bioenergetics are also dependent on Ca^2+^ signals, we assessed mitochondrial function by measuring the mitochondrial membrane potential (ΔΨm) using a membrane potential sensitive probe and the OCR using Seahorse technology. Our results show that in the SS mouse model, ΔΨm, which is critical for ATP synthesis, is diminished (Figure 10C and supplementary Figure 4). While the ATP-linked OCR remained unchanged, both the basal and maximal OCR were reduced. This suggests that mitochondrial functionality is compromised in the disease model, indicating a decreased capacity to respond to additional cellular stress. An intriguing question arises from these findings: are defects in the function of mitochondria a primary cause of fluid secretion loss in SS, or alternatively is this a consequence of disrupted [Ca^2+^]_i_ regulation? Moreover, DNA from damaged mitochondria can activate the cGAS/STING pathway, leading to inflammation(18,21). This implies that compromised mitochondria in early SS stages could trigger prolonged inflammation through the STING pathway, potentially contributing to SS progression. Understanding these mechanisms is crucial for developing effective treatments to halt or slow the progression of SS.

## Material and Methods

All animal procedures were approved by the University of Rochester Committee on Animal Resources (UCAR-2001-214E)

### Animals

The murine model of Sjögren’s syndrome was established through the induction of the STING pathway (56). Briefly, 8-10 weeks old female C57BL/6J wild type (WT) mice (Jackson Laboratory; Jax 000664) received subcutaneous injections of DMXAA (Vadimezan; GC16280) at a concentration of 25 mg/kg of body weight on both day 0 and day 21 of the experimental timeline (see figure 1). The control mouse received vehicle (5% sodium bicarbonate; Sigma-Aldrich; S8761), the DMXAA solvent at the corresponding time points. Experiments were performed on day 28 of the experimental timeline.

### Evaluation of saliva production

The mice were fasted for two hours prior to the evaluation of saliva production. The mice were anesthetized with a solution containing Ketamine (10 mg/mL) and Xylazine (1 mg/ml) by intraperitoneal injection (IP) at a dose of 7 μl/gm body weight over 2 minutes. The mouse was placed on a heating pad at 37℃ during experimentation. A Salimetrics Childen’s swab (Salimetrics; Cat. no. 5001.05) was placed within the oral cavity of each mouse. The mice were administered the muscarinic agonist pilocarpine (0.375 mg/kg body weight; Millipore Sigma; P6503) by IP injection. Two minutes after the pilocarpine injection, saliva was collected for the following 15 minutes. The saliva absorbed was subsequently separated from the moist swab through centrifugation at 10,000 rpm for 1 minute. The measurement of saliva weight served as a quantitative evaluation of the efficacy of whole saliva secretion. To measure neurotransmitter-stimulated saliva secretion more directly, the mouse was anesthetized as previously described (24) and a surgical incision was made in the skin to expose the submandibular gland (SMG). The surrounding connective tissue was excised to facilitate positioning within a custom-made 3D-printed gland holder. A pair of stimulation electrodes were attached to the duct bundle and the SMG. The pre-weighed filter paper was positioned within the oral cavity of the mouse to capture saliva secretion. Secretion was initiated by electrical stimulation sequences generated by a stimulus isolator (Iso-flex, A.M.P.I.) set at 5 mA, 200 ms, at frequencies of 1, 3, 5, 7, and 10 Hz with train frequency and duration (typically 1 minute) controlled by a train generator (DG2A, Warner Instruments). The interval between each stimulus was 3 minutes. After stimulation, the filter paper was removed and weighed. The difference between the weight of filter paper before and after the electrode stimulation represented the saliva produced by the respective salivary gland during the given stimulation period.

### *In vivo* Ca^2+^ imaging

Mist1^CreERT2+/–^ x GCaMP6f^+/–^ transgenic mice served as the experimental subjects for Ca^2+^ imaging of the submandibular gland (SMG) *in vivo*. The generation of Mist1^CreERT2+/–^ x GCaMP6f^+/–^transgenic mice by crossing GcaMP6f^flox^ mice (Jackson Laboratory; Jax 028865) with Mist1^CreERT2^ (Jackson Laboratory; Jax 029228, a gift from Dr. Catherine Ovitt, University of Rochester). A week before the DMXAA or 5% sodium bicarbonate injections, tamoxifen (Sigma-Aldrich; T5648) was given to the mice *via* oral gavage at a dose of 0.25 mg/g of body weight for 3 consecutive days to excise the loxP sites flanking the STOP codon allowing expression of the Ca^2+^ indicator within salivary glands. The mice were anesthetized and gland-exposed, as described previously (24,57). The immobilized gland was secured within the holder using a cover glass and maintained in Hank’s salt solution (HBSS). Ca^2+^ imaging was conducted *in vivo* via two-photon microscopy using an Olympus FVMPE-RS system equipped with an Insight X3 pulsed laser (Spectra-Physics) utilizing a heated (OKOLab COL2532) 25x water immersion lens (Olympus XLPlan N 1.05 W MP). GCaMP6f was excited at 950 nm and emission collected between 495–540 nm, with images captured at 0.5-second intervals following stimulation for 10 seconds with 3 minutes between stimulation periods. Statistical analyses were performed with two-way ANOVA with multiple comparisons using Prism (GraphPad) as indicated in the figure legends.

### Immunofluorescent staining for sliced tissue

Following verification of decreased saliva secretion in mice, glands were processed for immunocytochemistry. Briefly, the isolated salivary glands were fixed in 4% paraformaldehyde at 4℃ overnight. The fixed gland was processed, embedded in paraffin, and subsequently sliced into 5 μm thick sections. Two temperature-induced antigen retrieval protocols were used either based on HIER buffer (10 mM Tris-base, 1 mM EDTA-dehydrate, pH 9.2) or sodium citrate buffer (10 mM sodium citrate, 0.05% Tween 20, pH 6.0). Gland sections were blocked with the 10% donkey serum in 0.2% PBSA (PBS+ BSA) at room temperature (RT) for 1 hour. Sections were incubated with the primary antibody at 4℃ overnight (TMEM16a (Millipore Sigma; P6593; 1:250), Na^+^/K^+^ ATPase (Abcam; ab2872; 1:250), ATP5A (Abcam; ab14748; 1:500), AQP5 (Abcam; ab239904; 1:500), STING (Cell signaling Technology, Cat. 13647; 1:500)). Following washing, the sections were then incubated with the secondary antibody at RT for 1 hour (Donkey anti-rabbit Alexa 488 (Thermo Fisher Scientific; A-21206; 1:500), Donkey anti-mouse Alexa 594 (ThermoFisher Scientific; A-21203; 1:500)). Nuclei were identified by incubation in DAPI (Thermo Scientific^TM^; Cat. 62248; 1:1000) at RT for 5 minutes. Tissue sections were mounted using Immu-Mount solution on a slide and then sealed under a coverslip. Images were acquired by Olympus FV1000MP confocal microscopy employing an Olympus UPlanSApo 60x oil immersion objective. The analysis of images was performed using FIJI software. Statistical analyses were performed with a t-test using Prism (GraphPad) as indicated in the figure legends.

### Patch clamp electrophysiology

Acinar cells were allowed to adhere to Cell-Tak-coated glass coverslips for 15 minutes before experimentation. Coverslips were transferred to a chamber containing extracellular bath solution (155 mM tetraethylammonium chloride to block K^+^ channels, 2 mM CaCl_2_, 1 mM MgCl_2_, 10 mM HEPES, pH 7.2). Cl^-^ currents in individual cells were measured in the whole cell patch clamp configuration using pClamp 9 and an Axopatch 200B amplifier (Molecular Devices). Recordings were sampled at 2 kHz and filtered at 1 kHz. Pipette resistances were 3–5 MΩ, and seal resistances were greater than 1 GΩ. Pipette solutions (pH 7.2) contained 60 mM tetraethylammonium chloride, 90 mM tetraethylammonium glutamate, 10 mM HEPES, 1 mM HEDTA (N-(2-hydroxyethyl) ethylenediamine-N, N’, N’-triacetic acid) and 20 μM CaCl_2_ were used to mimic physiological buffering and basal [Ca^2+^]_i_ conditions (∼100 nM Ca^2+^). Free [Ca^2+^] was estimated using Maxchelator freeware. Agonists were directly perfused onto individual cells using a multibarrel perfusion pipette. The pipette solution for the increased basal [Ca^2+^]_i_ contained hEDTA and a free [Ca^2+^]_i_ of 500 nM, 1 μM, or 5 μM to induce calcium-activated Cl^-^ currents without the addition of any agonists. Experiments comparing EGTA, BAPTA, and HEDTA effects upon chloride currents induced by CCh stimulation contained 5 mM free concentrations of the chelator and 100 nM free [Ca^2+^] in the patch pipette. Chloride currents following agonist application were monitored with a single voltage step to 80 mV from a holding potential of −80 mV every second until current magnitudes reached a plateau. Current-voltage relationships were obtained by 20 mV incremental steps between −80 mv and 120 mV from a holding potential of −50 mV.

### STED microscopy

3D STED microscopy was performed using an Abberior Instruments Expert Line STED microscope equipped with an Olympus UPLSAPO ×100/1.4NA oil immersion objective. Briefly, lobules <1 mm were isolated following injection of saline beneath the capsule with a 29-gauge needle. The connecting tissue was digested in 0.1 mg/ml collagenase containing image buffer at 37℃ for 5 minutes. Then isolated lobules were fixed in 100% methanone at −20℃ for 5 minutes, and subsequently were blocked with 10% BSA in 0.1% PBST (PBS+ 0.1% Tween20) at RT for 1 hour with gentle shaking. Isolated lobules were incubated with primary antibodies overnight at 4°C (TMEM16a (Millipore Sigma; P6593; 1:300), IP_3_R3 (BD Transduction Laboratory; Cat. 610313; 1:200)). After being washed with 0.1% PBST, the membranes were incubated with secondary antibodies at RT for 1 hour (STAR RED, goat anti-rabbit IgG secondary antibody (Abberior, Cat#STRED-1001-500UG; 1:1000), Alexa Fluor 594 anti-rabbit IgG secondary antibody (Molecular Probes Cat#A-11037; 1:1000)). The tissue was mounted on the slides with Prolong^TM^ Gold antifade reagent (Invitrogen; Cat. P36930. Sequential confocal and STED images were obtained following excitation of Alexa Fluor 594 and STAR RED by 594 and 640 nm lasers, respectively. Both fluorophores were depleted in three dimensions with a 775 nm pulsed STED laser. Z-stacks were obtained by collecting images at 50 nm intervals using the 3D STED mode. Rescue STED was employed to minimize the light dosage. Blend mode depth projection images were generated and fluorophore volumes and interfaces between these volumes were analyzed using FIJI.

### Seahorse XF cell mito stress assay

Isolated SMGs were finely minced and subsequently resuspended in a solution composed of 0.5% Bovine Serum Albumin (BSA) in Hank’s Balanced Salt Solution (HBSS). To isolate acinar cells, the minced tissue was incubated in 0.5% BSA/HBSS containing 0.2 mg/ml of collagenase type II (Worthington; LS004204) for 30 minutes. Following this incubation, the suspension of cells was centrifuged at 500 rpm for 1 minute and the cellular pellet was then resuspended in 40 μg/ml of Trypsin inhibitor (Millipore; Cat. 65035) to terminate further digestion. The function of mitochondria was assessed in isolated acinar cells by measurement of oxygen consumption rate (OCR) employing a Seahorse XF Cell Mito Stress Test system (Agilent, USA). Briefly, Equal sized SMG cell pellets were suspended in buffer and 10 μl of the acinar cell suspension was seeded into individual wells of Seahorse cell culture microplates coated with 10 uL of Cell-Tak (0.25mg/ml) and the OCR was determined utilizing the Seahorse XFe96 extracellular flux analyzer following sequential exposure to 4μg/ml oligomycin (Millipore Sigma; O4876), 4µM carbonyl cyanide-4 (trifluoromethoxy)phenylhydrazone (FCCP; Millipore Sigma; C2920), and 0.5 µM rotenone/antimycin (Millipore Sigma; R8875; A8674) to measure the quantification of basal respiration, ATP-linked respiration, and maximum respiration rate, respectively. Statistical analyses were performed with, t-test using Prism (GraphPad) as indicated in the figure legends.

### Measurement of mitochondrial membrane potential

Isolated SMG acinar cells were loaded with 20 nM Tetramethylrhodamine, Ethyl Ester (TMRE; ThermoFisher Scientific: T669), and 1μM of MitoTracker Green (Invitrogen^TM^; M7514). Fluorescence of both TMRE and MitoTracker Green was captured simultaneously using an inverted epifluorescence Nikon microscope with a 40x oil immersion objective. The TMRE fluorescence was excited at 560 nm and emitted light collected at 574 nm; MitoTracker Green was excited at 488 nm and emitted light collected at 530 nm. Images were obtained every 1 s with an exposure of 20 ms and 4 x 4 binning using a digital camera controlled by TILL Photonics, TILLvision software. The acinar cells were exposed to 4 μM FCCP for 3 minutes by perfusion to rapidly dissipate the membrane potential. Mitochondrial membrane potential was quantified as the change in the ratio of TMRE/MitoTracker Green fluorescence before and after the administration of FCCP. Statistical analyses were performed with a t-test using Prism (GraphPad) as indicated in the figure legends.

### Western blotting

Finely minced salivary glands were homogenized in a lysis buffer supplemented with protease inhibitor cocktail (Complete mini; Roche Diagnostics) for 16-20 strokes. After incubating on ice for 30 minutes, solubilized proteins were separated by centrifugation at 13000 rpm at 4℃ for 30 minutes. 10μg of protein lysate was loaded on 7.5%-12% SDS-polyacrylamide gels. Subsequently, the proteins were transferred to PVDF membranes at a voltage of 35V at 4°C overnight. The membrane was blocked with 5% non-fat skimmed milk in TBST (50 mM Tris-HCl, pH 7.5 with 0.1% Tween20) at RT for 1 hour and subsequently incubated with primary antibodies overnight at 4°C (Actin (Millipore Sigma; A2228; 1:10000), IP_3_R2 (Antibody Research Corporation; 1:1000), IP_3_R3 (BD Transduction Laboratory; Cat. 610313; 1:1000), TMEM16a (Abcam; ab84115; 1:1000)). After being washed with 0.1% TBST, the membranes were incubated with secondary antibodies at RT for 1 hour (Goat anti-rabbit IgG (H&L) (Invitrogen; SA535571; 1:10000), Goat anti-mouse IgG (H&L) (Invitrogen; SA535521; 1:10000)). Protein band intensity from western blotting was quantified by FIJI. The relative ratio of DMXAA-treated/ vehicle control was calculated in Excel. Lastly, graphical generation and statistics were performed with a t-test using Prism (GraphPad) as indicated in the figure legends.

## Acknowledgments

The authors gratefully acknowledge the University of Rochester’s Center for Advanced Microscopy and Nanoscopy (CALMN) for providing access to Multiphoton microscopy for in vivo live imaging and STED super-resolution microscopy, and for Center for Advanced Research Technologies (CART) for the Electron & cryo Microscopy Resource. We also thank the Flow Cytometry Resource (FCR) for its support with the mitochondrial stress assay. Special thanks to Dr. Paul Brooks for sharing Seahorse Technology XF analyzers and for engaging in discussions for optimization of experiments. Thanks to Dr. Catherine Ovitt for her instruction on tissue staining techniques. Additionally, we wish to express our appreciation to all members of the Yule laboratory for their invaluable feedback, discussions, and assistance, which have been essential in advancing this study. The work was supported by a grant from NIH (NIDCR) DE014756 (to DIY).

---

**Supplementary Figure 1.**
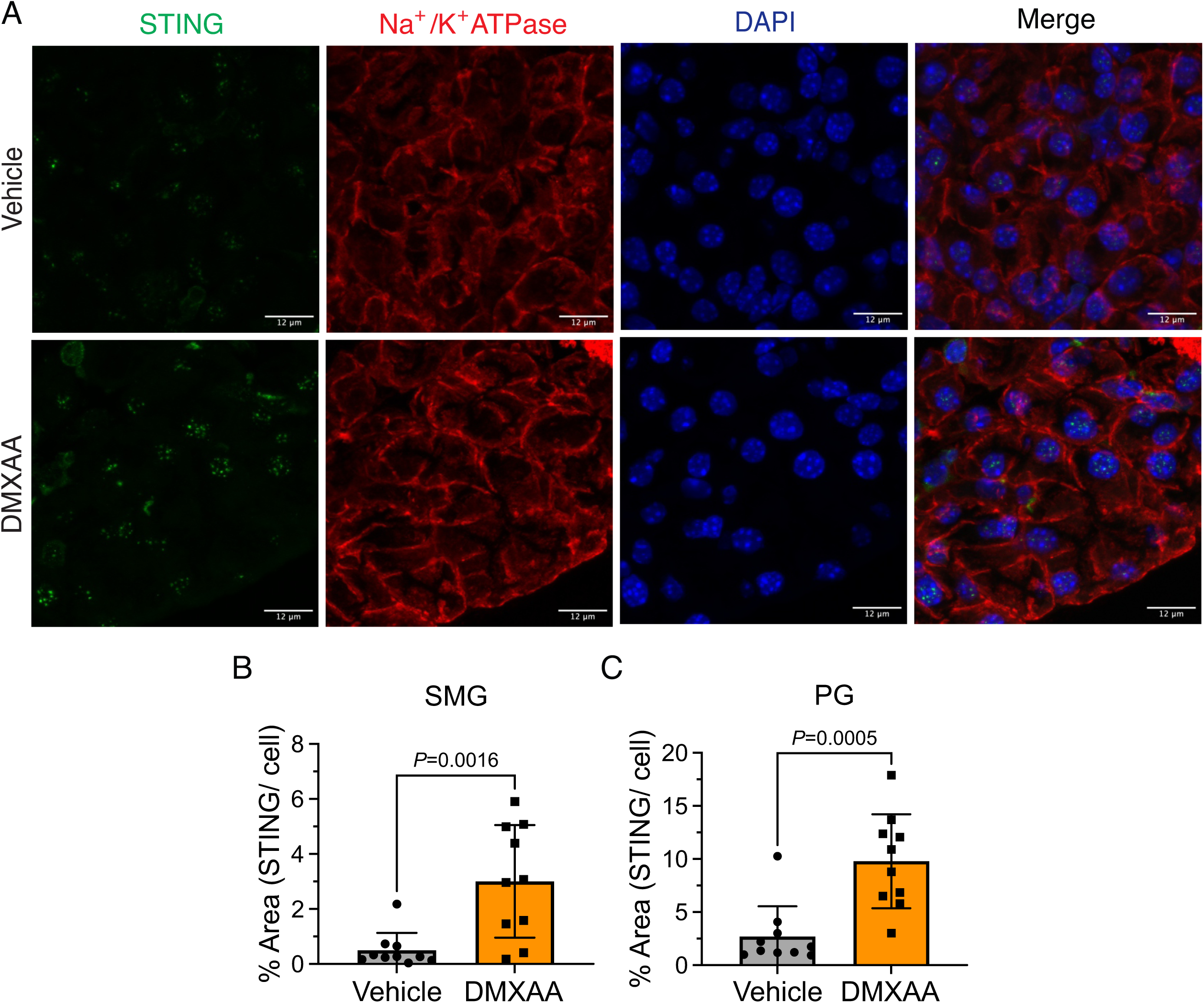
Up-regulation of STING protein expression in both SMG and PG treated with DMXAA. Immunofluorescent staining in SMG tissue for STING (green), Na^+^/K^+^ ATPase (red), and DAPI for nucleus (blue). The upper panel is in vehicle-treated condition and the bottom panel is the SS mouse model. Scale bar: 12 μm. (B-C) STING protein expression was quantified by the percentage of a cell occupied by STING protein in (B) SMG and (C) PG. Vehicle, N= 3 mice; SS mouse model: N= 3 mice. Unpaired two-tailed t-test.

**Supplementary Figure 2.**
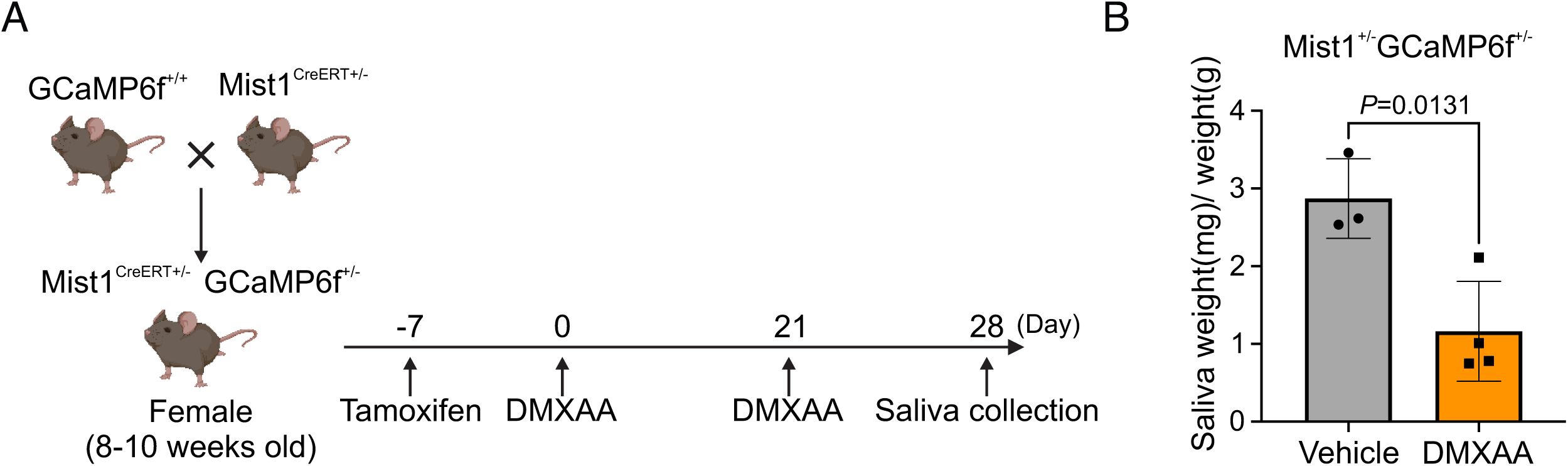
Deficiency in secretion in Mist1^CreERT+/-^GCaMP6f^+/-^ genetic mouse treated with DMXAA. (A) Schematic timeline for the generation of the SS mouse model in the Mist1^CreERT+/-^GCaMP6f^+/-^ genetic mouse. The female Mist1^CreERT+/-^GcaMP6f^+/-^ mouse received two subcutaneous doses of DMXAA on day 0 and day 21. The salivary gland function was assessed on day 28. (B) The gland function was evaluated by the weight of pilocarpine-induced saliva, normalized to each mouse’s weight. Vehicle = 3 mice, DMXAA= 4 mice. Mean ± SD. Unpaired two-tailed t-test.

**Supplementary Figure 3.**
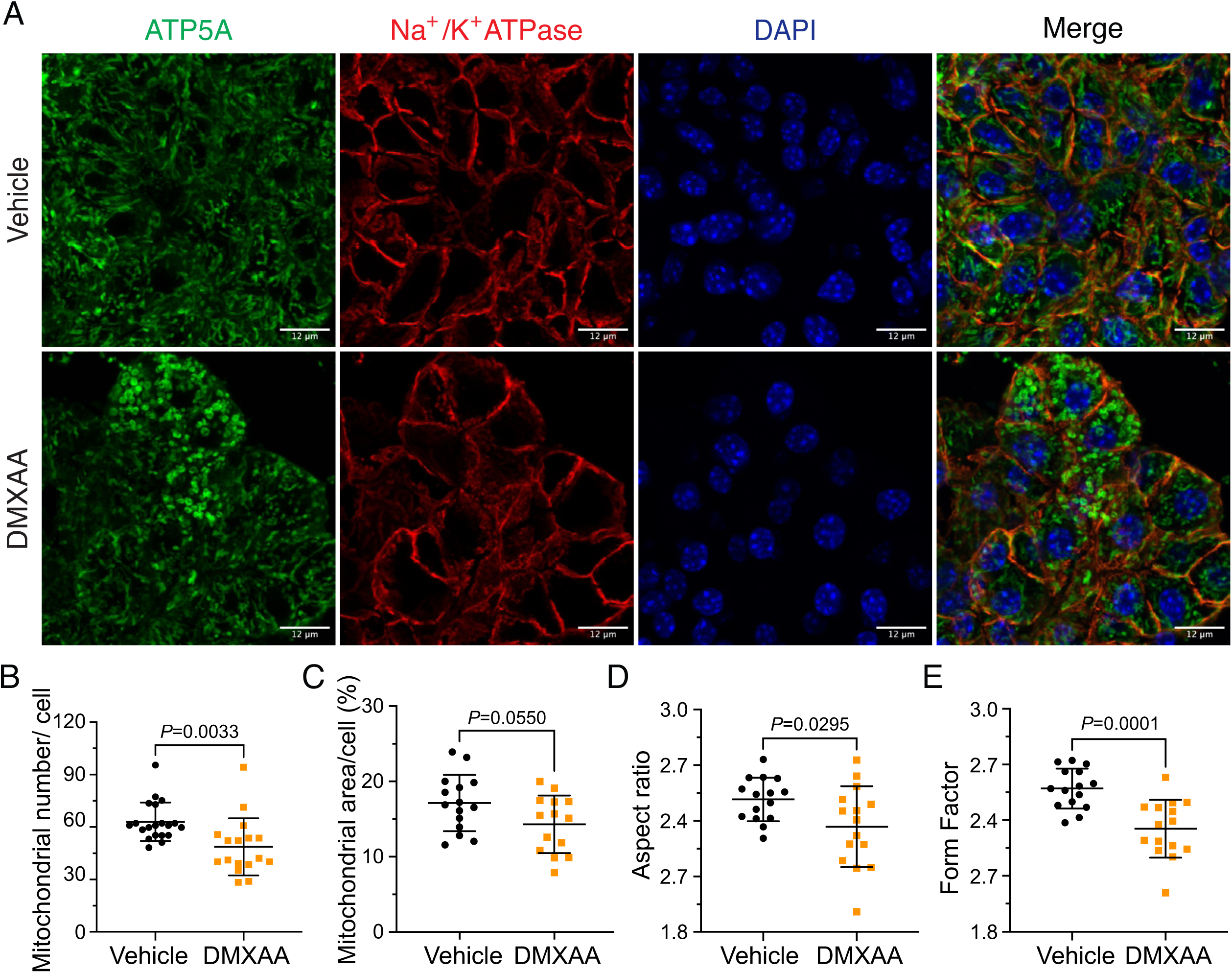
Mitochondrial alterations in the parotid gland of SS mouse model. (A) Immunofluorescent staining in PG tissue for ATP5A (green), Na^+^/K^+^ ATPase (red), and DAPI for nucleus (blue). The upper panel is from vehicle-treated animals and the bottom panel is from the SS mouse model. Scale bar: 12 μm. The mitochondrial content was quantified by (B) the mitochondrial number per acinar cell and (C) the percentage of area occupied by mitochondria per acinar cell. The mitochondrial morphology was analyzed by the (D) AR for the degree of mitochondrial tubular shape and (E) FF for the degree of mitochondrial branching (complexity). In (B) to (E), black dots represent the vehicle condition, and orange squares indicate the SS mouse model. Each symbol represents the mean of 10 cells per image. Vehicle: N= 20 and SS mouse model: N= 15-17 from four mice. Mean ± SD. Unpaired two-tailed t-test.

**Supplementary Figure 4.**
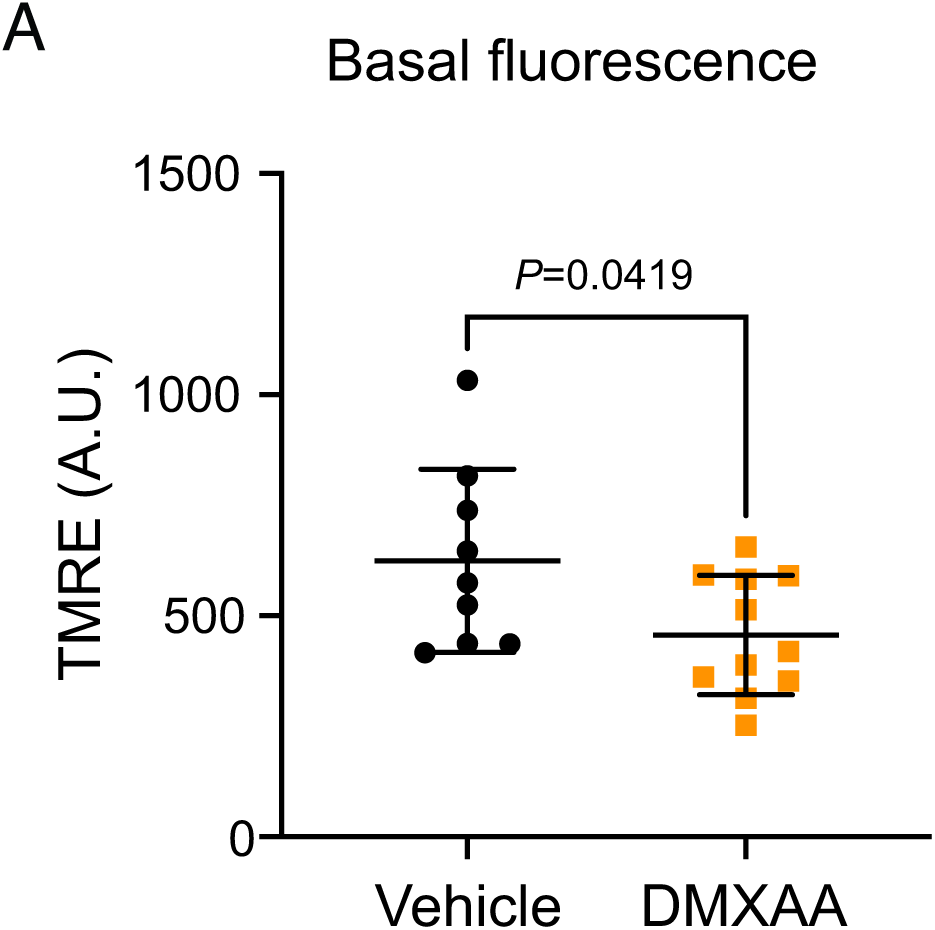
Reduction of the basal TMRE fluorescence in DMXAA-treated animals. (A) The fluorescent intensity of TMRE loading in isolated acinar cells from vehicle-treated and DMXAA-treated mice. Black dots represent the vehicle condition, and orange squares indicate the SS mouse model. Each symbol represents the mean of 10 cells per image. Vehicle: N= 9 and SS mouse model: N= 11 from three mice. Mean ± SD. Unpaired two-tailed t-test.

**Supplementary Figure 5.**
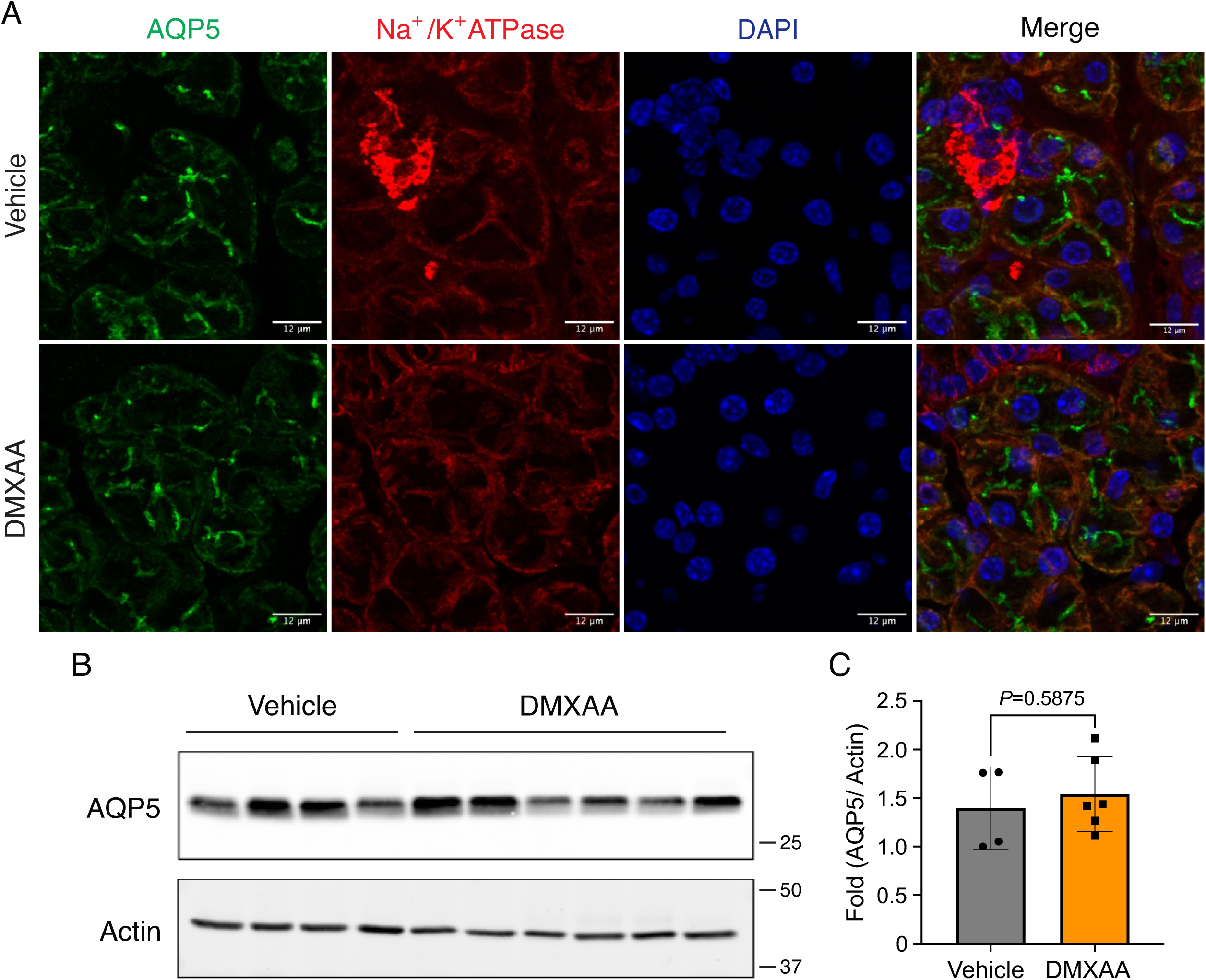
AQP5, the water channel, remained the comparable expression and proper localization in SMG in the disease mouse model. (A) Immunofluorescent staining in SMG tissue for AQP5 (green), Na^+^/K^+^ ATPase (red), and DAPI for nucleus (blue). The upper panel is from vehicle condition and the bottom panel is from the SS mouse model. Scale bar: 12 μm. (B) Western blotting showed the protein level of AQP5 in SMG from vehicle-treated control and disease mouse model. (C) The quantification of AQP5 normalized to internal control, Actin. Vehicle: N= 4; SS mouse model: N= 6. Mean ± SD. Unpaired two-tailed t-test.

**Supplementary Figure 6.**
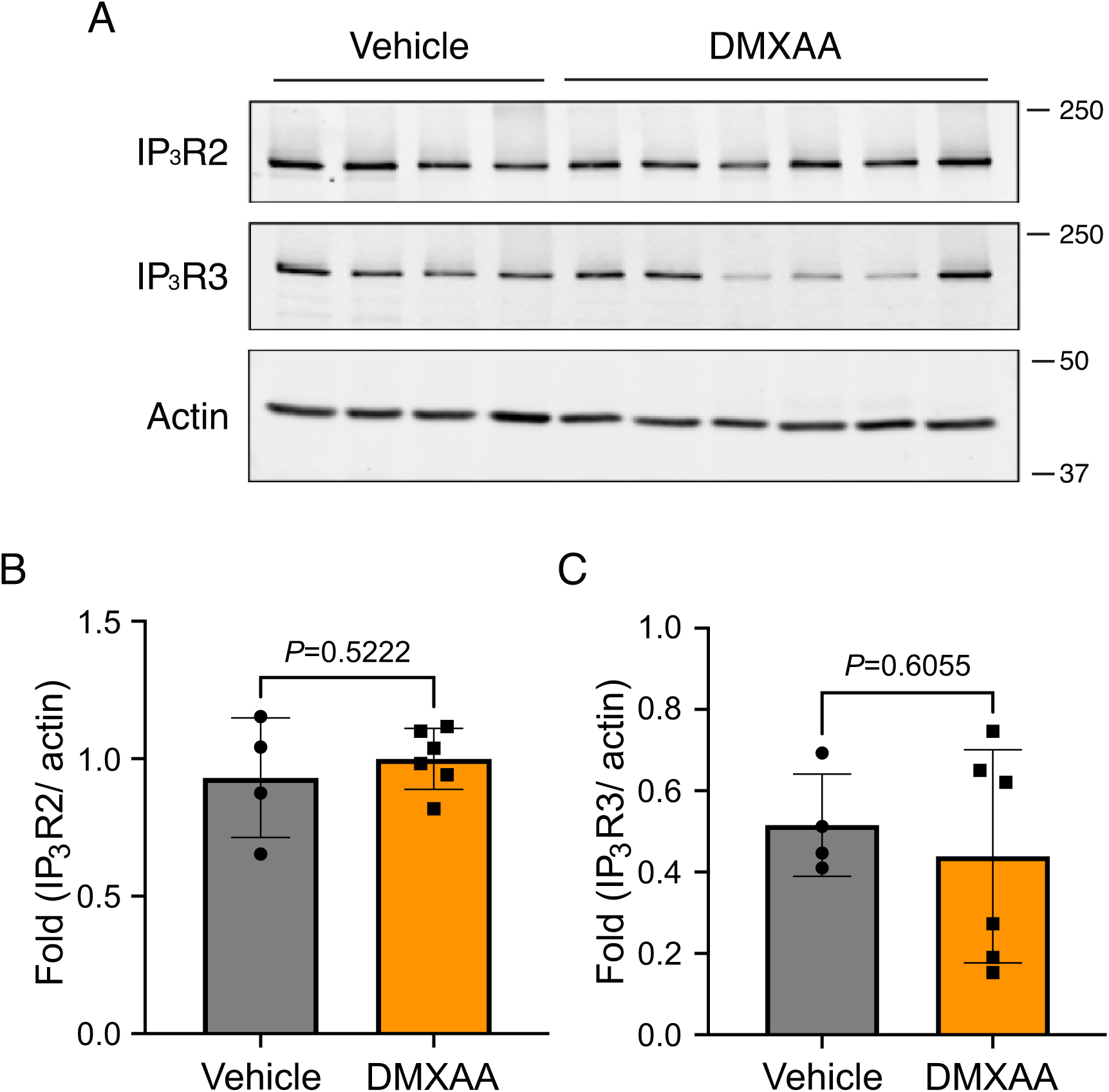
No significant alteration in IP_3_R protein levels in SMG in the DMXAA-treated mouse model. (A) Western blotting indicated the IP_3_R2 and IP_3_R3 protein levels from vehicle-treated control and SS mouse models. Actin was probed as the internal control. (B) The quantification of IP_3_R2 and IP_3_R3 normalized to Actin, the internal control. Vehicle: N= 4; SS mouse model: N= 6. Mean ± SD. Unpaired two-tailed t-test.

## References

1. Williamson, S. P. H. a. R. T. (2001) <A review of saliva-Normal composition, flow, and function.pdf>. J Prosthet Dent 85

2. Carpenter, G. H. (2013) The secretion, components, and properties of saliva. Annu Rev Food Sci Technol 4, 267–276

3. Pedersen, A. M. L., Sorensen, C. E., Proctor, G. B., Carpenter, G. H., and Ekstrom, J. (2018) Salivary secretion in health and disease. J Oral Rehabil 45, 730–746

4. de Paula, F., Teshima, T. H. N., Hsieh, R., Souza, M. M., Nico, M. M. S., and Lourenco, S. V. (2017) Overview of Human Salivary Glands: Highlights of Morphology and Developing Processes. Anat Rec (Hoboken) 300, 1180–1188

5. Futatsugi, A., Nakamura, T., Yamada, M. K., Ebisui, E., Nakamura, K., Uchida, K., Kitaguchi, T., Takahashi-Iwanaga, H., Noda, T., Aruga, J., and Mikoshiba, K. (2005) IP3 receptor types 2 and 3 mediate exocrine secretion underlying energy metabolism. Science 309, 2232–2234

6. Lee, M. G., Xu, X., Zeng, W., Diaz, J., Wojcikiewicz, R. J., Kuo, T. H., Wuytack, F., Racymaekers, L., and Muallem, S. (1997) Polarized expression of Ca2+ channels in pancreatic and salivary gland cells. J Biol Chem 272, 15765–15770

7. Pages, N., Vera-Siguenza, E., Rugis, J., Kirk, V., Yule, D. I., and Sneyd, J. (2019) A Model of Ca2+ Dynamics in an Accurate Reconstruction of Parotid Acinar Cells. Bull Math Biol 81, 1394–1426

8. Melvin, J. E., Yule, D., Shuttleworth, T., and Begenisich, T. (2005) Regulation of fluid and electrolyte secretion in salivary gland acinar cells. Annu Rev Physiol 67, 445–469

9. Saleh, J., Figueiredo, M. A., Cherubini, K., and Salum, F. G. (2015) Salivary hypofunction: an update on aetiology, diagnosis and therapeutics. Arch Oral Biol 60, 242–255

10. Khalid F. Tabbara, M., and Carlos L. Vera-Cristo, MD. (2000) Sjögren syndrome. Current Opinion in Ophthalmology 11, 449–454

11. Ramos-Casals, M., Brito-Zeron, P., Siso-Almirall, A., and Bosch, X. (2012) Primary Sjogren syndrome. BMJ 344, e3821

12. Clio P. Mavragani MD, H. M. M. M. (2014) Sjögren syndrome. Canadian Medical Association Journal 10

13. Brito-Zeron, P., Baldini, C., Bootsma, H., Bowman, S. J., Jonsson, R., Mariette, X., Sivils, K., Theander, E., Tzioufas, A., and Ramos-Casals, M. (2016) Sjogren syndrome. Nat Rev Dis Primers 2, 16047

14. Kiripolsky, J., Shen, L., Liang, Y., Li, A., Suresh, L., Lian, Y., Li, Q. Z., Gaile, D. P., and Kramer, J. M. (2017) Systemic manifestations of primary Sjogren’s syndrome in the NOD.B10Sn-H2(b)/J mouse model. Clin Immunol 183, 225–232

15. Jonsson, R., Brokstad, K. A., Jonsson, M. V., Delaleu, N., and Skarstein, K. (2018) Current concepts on Sjogren’s syndrome - classification criteria and biomarkers. Eur J Oral Sci 126 Suppl 1, 37–48

16. Lee, B. H., Gauna, A. E., Pauley, K. M., Park, Y. J., and Cha, S. (2012) Animal models in autoimmune diseases: lessons learned from mouse models for Sjogren’s syndrome. Clin Rev Allergy Immunol 42, 35–44

17. Gao, Y., Chen, Y., Zhang, Z., Yu, X., and Zheng, J. (2020) Recent Advances in Mouse Models of Sjogren’s Syndrome. Front Immunol 11, 1158

18. Decout, A., Katz, J. D., Venkatraman, S., and Ablasser, A. (2021) The cGAS-STING pathway as a therapeutic target in inflammatory diseases. Nat Rev Immunol 21, 548–569

19. J. Papinska, H. B., G.B. Gmyrek, M. Sroka, S. Tummala, K.A. Fitzgerald, and U.S. Deshmukh. (2018) Activation of Stimulator of Interferon Genes (STING) and Sjögren Syndrome. Journal of Dental Research 97, 893–900

20. Huijser, E., Bodewes, I. L. A., Lourens, M. S., van Helden-Meeuwsen, C. G., van den Bosch, T. P. P., Grashof, D. G. B., van de Werken, H. J. G., Lopes, A. P., van Roon, J. A. G., van Daele, P. L. A., Brkic, Z., Dik, W. A., and Versnel, M. A. (2022) Hyperresponsive cytosolic DNA-sensing pathway in monocytes from primary Sjogren’s syndrome. Rheumatology (Oxford) 61, 3491–3496

21. Gao, P., Ascano, M., Zillinger, T., Wang, W., Dai, P., Serganov, A. A., Gaffney, B. L., Shuman, S., Jones, R. A., Deng, L., Hartmann, G., Barchet, W., Tuschl, T., and Patel, D. J. (2013) Structure-function analysis of STING activation by c[G(2’,5’)pA(3’,5’)p] and targeting by antiviral DMXAA. Cell 154, 748–762

22. Weiss, J. M., Guerin, M. V., Regnier, F., Renault, G., Galy-Fauroux, I., Vimeux, L., Feuillet, V., Peranzoni, E., Thoreau, M., Trautmann, A., and Bercovici, N. (2017) The STING agonist DMXAA triggers a cooperation between T lymphocytes and myeloid cells that leads to tumor regression. Oncoimmunology 6, e1346765

23. Ceron, S., North, B. J., Taylor, S. A., and Leib, D. A. (2019) The STING agonist 5,6-dimethylxanthenone-4-acetic acid (DMXAA) stimulates an antiviral state and protects mice against herpes simplex virus-induced neurological disease. Virology 529, 23–28

24. Takano, T., Wahl, A. M., Huang, K. T., Narita, T., Rugis, J., Sneyd, J., and Yule, D. I. (2021) Highly localized intracellular Ca(2+) signals promote optimal salivary gland fluid secretion. Elife 10

25. Eisner, D., Neher, E., Taschenberger, H., and Smith, G. (2023) Physiology of intracellular calcium buffering. Physiol Rev 103, 2767–2845

26. Duchen, M. R. (2000) Mitochondria and calcium-from cell signalling to cell death. Journal of Physiology 529, 57–68

27. Csordas, G., Renken, C., Varnai, P., Walter, L., Weaver, D., Buttle, K. F., Balla, T., Mannella, C. A., and Hajnoczky, G. (2006) Structural and functional features and significance of the physical linkage between ER and mitochondria. J Cell Biol 174, 915–921

28. Ye, L., Zeng, Q., Ling, M., Ma, R., Chen, H., Lin, F., Li, Z., and Pan, L. (2021) Inhibition of IP3R/Ca2+ Dysregulation Protects Mice From Ventilator-Induced Lung Injury via Endoplasmic Reticulum and Mitochondrial Pathways. Front Immunol 12, 729094

29. Katona, M., Bartok, A., Nichtova, Z., Csordas, G., Berezhnaya, E., Weaver, D., Ghosh, A., Varnai, P., Yule, D. I., and Hajnoczky, G. (2022) Capture at the ER-mitochondrial contacts licenses IP(3) receptors to stimulate local Ca(2+) transfer and oxidative metabolism. Nat Commun 13, 6779

30. Barrera, M. J., Aguilera, S., Castro, I., Carvajal, P., Jara, D., Molina, C., Gonzalez, S., and Gonzalez, M. J. (2021) Dysfunctional mitochondria as critical players in the inflammation of autoimmune diseases: Potential role in Sjogren’s syndrome. Autoimmun Rev 20, 102867

31. Valente, A. J., Maddalena, L. A., Robb, E. L., Moradi, F., and Stuart, J. A. (2017) A simple ImageJ macro tool for analyzing mitochondrial network morphology in mammalian cell culture. Acta Histochem 119, 315–326

32. Harwig, M. C., Viana, M. P., Egner, J. M., Harwig, J. J., Widlansky, M. E., Rafelski, S. M., and Hill, R. B. (2018) Methods for imaging mammalian mitochondrial morphology: A prospective on MitoGraph. Anal Biochem 552, 81–99

33. Hogan, M. C., Stary, C. M., Balaban, R. S., and Combs, C. A. (2005) NAD(P)H fluorescence imaging of mitochondrial metabolism in contracting Xenopus skeletal muscle fibers: effect of oxygen availability. J Appl Physiol (1985) 98, 1420–1426

34. Martinez, J., Marmisolle, I., Tarallo, D., and Quijano, C. (2020) Mitochondrial Bioenergetics and Dynamics in Secretion Processes. Front Endocrinol (Lausanne) 11, 319

35. Navaratnarajah, T., Anand, R., Reichert, A. S., and Distelmaier, F. (2021) The relevance of mitochondrial morphology for human disease. Int J Biochem Cell Biol 134, 105951

36. Schaefer, P. M., Kalinina, S., Rueck, A., von Arnim, C. A. F., and von Einem, B. (2019) NADH Autofluorescence-A Marker on its Way to Boost Bioenergetic Research. Cytometry A 95, 34–46

37. Zorova, L. D., Popkov, V. A., Plotnikov, E. Y., Silachev, D. N., Pevzner, I. B., Jankauskas, S. S., Babenko, V. A., Zorov, S. D., Balakireva, A. V., Juhaszova, M., Sollott, S. J., and Zorov, D. B. (2018) Mitochondrial membrane potential. Anal Biochem 552, 50–59

38. Yanfei Hu, Y. N., Karnam R. Purushotham, and Michael G. Humphreys-Beher. (1992) Functional changes in salivary glands of autoimmune disease-prone NOD mice. Journal of Physiology

39. Shen, L., Zhang, C., Wang, T., Brooks, S., Ford, R. J., Lin-Lee, Y. C., Kasianowicz, A., Kumar, V., Martin, L., Liang, P., Cowell, J., and Ambrus, J. L., Jr. (2006) Development of autoimmunity in IL-14alpha-transgenic mice. J Immunol 177, 5676–5686

40. Shen, L., Suresh, L., Li, H., Zhang, C., Kumar, V., Pankewycz, O., and Ambrus, J. L., Jr. (2009) IL-14 alpha, the nexus for primary Sjogren’s disease in mice and humans. Clin Immunol 130, 304–312

41. Papinska, J., Bagavant, H., Gmyrek, G. B., and Deshmukh, U. S. (2020) Pulmonary Involvement in a Mouse Model of Sjogren’s Syndrome Induced by STING Activation. Int J Mol Sci 21

42. Phillips, R. M. (1999) Inhibition of DT-diaphorase (NAD(P)H:quinone oxidoreductase, EC 1.6.99.2) by 5,6-dimethylxanthenone-4-acetic acid (DMXAA) and flavone-8-acetic acid (FAA): implications for bioreductive drug development. Biochem Pharmacol 58, 303–310

43. Teos, L. Y., Zhang, Y., Cotrim, A. P., Swaim, W., Won, J. H., Ambrus, J., Shen, L., Bebris, L., Grisius, M., Jang, S. I., Yule, D. I., Ambudkar, I. S., and Alevizos, I. (2015) IP3R deficit underlies loss of salivary fluid secretion in Sjogren’s Syndrome. Sci Rep 5, 13953

44. Jin, X., Shah, S., Du, X., Zhang, H., and Gamper, N. (2016) Activation of Ca(2+)-activated Cl(-) channel ANO1 by localized Ca(2+) signals. J Physiol 594, 19–30

45. Romanenko, V. G., Catalan, M. A., Brown, D. A., Putzier, I., Hartzell, H. C., Marmorstein, A. D., Gonzalez-Begne, M., Rock, J. R., Harfe, B. D., and Melvin, J. E. (2010) Tmem16A encodes the Ca2+-activated Cl-channel in mouse submandibular salivary gland acinar cells. J Biol Chem 285, 12990–13001

46. Tsubota, K., Hirai, S., King, L. S., Agre, P., and Ishida, N. (2001) Defective cellular trafficking of lacrimal gland aquaporin-5 in Sjogren’s syndrome. Lancet 357, 688–689

47. Catalan, M. A., Kondo, Y., Pena-Munzenmayer, G., Jaramillo, Y., Liu, F., Choi, S., Crandall, E., Borok, Z., Flodby, P., Shull, G. E., and Melvin, J. E. (2015) A fluid secretion pathway unmasked by acinar-specific Tmem16A gene ablation in the adult mouse salivary gland. Proc Natl Acad Sci U S A 112, 2263–2268

48. Zeng, M., Szymczak, M., Ahuja, M., Zheng, C., Yin, H., Swaim, W., Chiorini, J. A., Bridges, R. J., and Muallem, S. (2017) Restoration of CFTR Activity in Ducts Rescues Acinar Cell Function and Reduces Inflammation in Pancreatic and Salivary Glands of Mice. Gastroenterology 153, 1148–1159

49. Gresz, V., Horvath, A., Gera, I., Nielsen, S., and Zelles, T. (2015) Immunolocalization of AQP5 in resting and stimulated normal labial glands and in Sjogren’s syndrome. Oral Dis 21, e114–120

50. Shihab Shah, C. M. C., Pierce Mullen, Stephen Milne, Viktor Lukacs, Mark S. Shapiro, Nikita Gamper. (2020) Local Ca2+ signals couple activation of TRPV1 and ANO1 sensory ion channels. Science Signaling 13

51. Wang, Q., Bai, L., Luo, S., Wang, T., Yang, F., Xia, J., Wang, H., Ma, K., Liu, M., Wu, S., Wang, H., Guo, S., Sun, X., and Xiao, Q. (2020) TMEM16A Ca(2+)-activated Cl(-) channel inhibition ameliorates acute pancreatitis via the IP(3)R/Ca(2+)/NFkappaB/IL-6 signaling pathway. J Adv Res 23, 25–35

52. Denton, R. M. (2009) Regulation of mitochondrial dehydrogenases by calcium ions. Biochim Biophys Acta 1787, 1309–1316

53. Detmer, S. A., and Chan, D. C. (2007) Functions and dysfunctions of mitochondrial dynamics. Nat Rev Mol Cell Biol 8, 870–879

54. Porat-Shliom, N., Harding, O. J., Malec, L., Narayan, K., and Weigert, R. (2019) Mitochondrial Populations Exhibit Differential Dynamic Responses to Increased Energy Demand during Exocytosis In Vivo. iScience 11, 440–449

55. Li, N., Li, Y., Hu, J., Wu, Y., Yang, J., Fan, H., Li, L., Luo, D., Ye, Y., Gao, Y., Xu, H., Hai, W., and Jiang, L. (2022) A Link Between Mitochondrial Dysfunction and the Immune Microenvironment of Salivary Glands in Primary Sjogren’s Syndrome. Front Immunol 13, 845209

56. Bagavant, H., and Deshmukh, U. S. (2020) Protocols for Experimental Sjogren’s Syndrome. Curr Protoc Immunol 131, e114

57. Takano, T., and Yule, D. I. (2022) In vivo Ca(2+) Imaging in Mouse Salivary Glands. Bio Protoc 12, e4380

